# Isolation and screening of heavy metal resistant ammonia oxidizing bacteria from soil and waste dump: a potential candidates for bioremediation of heavy metals

**DOI:** 10.1101/836890

**Authors:** Veronica Fabian Nyoyoko, Chukwudi U Anyanwu

## Abstract

The study was undertaken to examine the response of ammonia oxidizing bacteria to different heavy metal salt in an elevated concentration. Surface soil samples at depth of 0-15 cm were collected at random from Akwa Ibom State University in Akwa Ibom State, soil sample from University of Nigeria, Nsukka and from solid waste disposal site in Uyo, Akwa Ibom State. The response of heavy metal salt on Ammonia Oxidizing bacteria(AOB) isolated from soil samples were investigated by supplementing different heavy metal salts namely, copper(Cu),nickel (Ni), lead (Pb) and cadmium (Cd) at four loading rates(100,200,500,1000 µg/ml) in mineral salt broth with Ammonia Oxidizing Bacteria (AOB) isolate. The cultures were incubated for 7 days. Growth of AOB was measured by withdrawing samples from the medium every 24 hours and absorbance of the turbidity measured at 600 nanometre using spectrophotometer. All bacteria showed high tendency to decrease optical density while increasing metal concentration in the medium. Tolerance for the metal ions was dependent on concentration, time and the isolate tested. All the Ammonia oxidizing bacterial (AOB) showed a high level of tolerance for the metals tested, and exhibited good growth at all metal salt concentrations tested. These make the nitrifying bacteria attractive potential candidates for further investigations regarding their ability to remove metals from contaminated soil.

**IMPORTANCE:** The aim of this study is tolerance of ammonia oxidizing bacteria growth to heavy metal. Nitrifying bacteria remain a good option for bioremediation of soil and waste dump, since it is regarded as eco-friendly and efficient in biosorption of heavy metal. The study is significant to the field of environmental microbiology by adding to knowledge in bioremediation.

## INTRODUCTION

Environmental pollution has been on the rise in the past few decades owing to increased human activities on energy reservoirs, unsafe agricultural practices and rapid industrialization (Hadia and Ahmed, 2018). Amongst the pollutants that are of environmental and public health concerns due to their toxicities are: heavy metals, nuclear wastes, pesticides, greenhouse gases, and hydrocarbons. These toxic pollutants are discharged from specific locations worldwide and thus pollute specific regions. These pollutants have different toxicity and chemical behaviour (Nibourg *et al.*, 2013). Environmental contamination caused by heavy metals (HM) has received increased attention worldwide (Gupta *et al.*, 2016; Sharma, 2016; Ayangbenro and Babalola, 2017; Liu *et al.*, 2018). Heavy metal (HM) is any metallic element that has a relatively high density and is toxic or poisonous at low concentrations. Heavy metals are elements with atomic number higher than 20, an atomic mass greater than 40 g and a specific weight of more than 5 g/cm^3^ (Canli *et al.*, 2003; Hamsa *et al.*, 2017). These elements often find their way into soil through environmental contaminants including the atmospheric pollutants in industrial regions (emissions from the rapidly expanding industrial areas), unlimited use of agricultural fertilizers, mine tailings, disposal of high metal wastes, leaded gasoline and paints, animal manures, sewage sludge, pesticides, wastewater irrigation, coal combustion residues, spillage of petrochemicals, atmospheric deposition, municipal and industrial sewage systems in a nonreturnable fashion (Cobbina *et al*., 2015; Malik *et al.*, 2017; Srivastava *et al.*, 2017; Ramya *et al.*, 2018).

Activities such as the use of agrochemicals and long-term application of urban sewage sludge, industrial waste disposal, waste incineration, and vehicle exhausts are the main sources of HM in agricultural soils (Mishra *et al.*, 2017). Heavy metals in the soil include mercury (Hg), lead (Pb), chromium (Cr), arsenic (As), zinc (Zn), cadmium (Cd), uranium (U), selenium (Se), silver (Ag), gold (Au), copper (Cu) and nickel (Ni) (Srivastava *et al.*, 2017). The danger of heavy metals is intensified by their almost indefinite persistence in the environment due to their absolute nature which cannot be degraded (Gupta *et al.*, 2016). Metals are non-biodegradable but can be transformed through sorption, methylation, complexation and changes in valence state (Anyanwu *et al.*, 2011).

Toxic metals apply their toxicity in the displacement of essential metals from their normal binding sites of biological molecules, inhibition of enzymatic functioning and disruption of nucleic acid structure, oxidation stress, genotoxicity and interfering with signalling pathways (Srivastava *et al.*, 2017; Venkatachalam *et al.*, 2017). Ecologically, the accumulation of heavy metals in soils is extremely hazardous because soil is a major link in the natural cycling of chemical elements; it is also a primary component of the trophic chain (Liu *et al.*, 2012; Sagi and Yigit, 2012; Wyszkowska, 2013).

Industrial operations such as electroplating, steel manufacturing, leather tanning, wood preservation, ceramics, glass manufacturing, chemical processing and fertilizer applications release alarmingly higher amounts of heavy metals into the natural environment (Ramya *et al.*, 2018). Pollution by heavy metal is a threat to the environment and it remediation is a major challenges to environmental research. Heavy metal pollution is a serious global environmental problem as it adversely affects biotic and abiotic components of the ecosystem and alters the composition and activity of soil microbial communities (Ayangbenro and Babalola, 2017). The non-biodegradability of heavy metals makes it hard to remove them from contaminated biological tissues and soil and this is a major concern for global health because of their lethal nature.

Nitrification is a biochemical process of oxidation of ammonia (NH_4_^+^) to nitrite (NO_2_^-^), then finally to nitrate (NO_3_^-^) by nitrifying bacteria. Nitrification is catalysed by two types of reactions. The first type of reaction is the oxidation of ammonia to nitrite by ammonium oxidizing bacteria (AOB). The second type of reaction involves the oxidation of nitrite to nitrate by nitrite-oxidizing bacteria (NOB) (Ma *et al*., 2014; Capone, 2018). Ammonium-oxidizing microorganisms are organism that carries out the first step in nitrification reaction (biochemical process of oxidation of ammonia (NH_4_^+^). They include ammonia oxidizing bacteria (AOB) *(Nitrosomonas, Nitrosococcus, Nitrosospira, Nitrosolobus, Nitrosovibrio)*, ammonia-oxidizing archaea (AOA), and heterotrophic bacteria (*Arthrobacter globiformis, Aerobacter aerogenes, Thiosphaera pantotropha, Streptomyces grisens*, and various *Pseudomonas* spp*) and* fungi (*Aspergillus flavus)* (Hamsa *et al.*, 2017). Recent research on the metabolic pathways of heterotrophic ammonia oxidation has been conducted using *Paracoccus denitrificans* (Moir *et al.*, 1996b), *Alcaligenes faecalis* (Joo *et al.*, 2005), *Pseudomonas putida* (Daum *et al.*, 1998), and a few other bacterial species. Some studies have suggested that the biochemical mechanisms of heterotrophic nitrification differ among strains (Niel *et al.*, 1990; Wehrfritz *et al.*, 1993). *Nitrospira* in the Nitrogen Oxidizing Bacteria group have been reported as complete ammonia oxidizing bacteria (comammox) that perform the complete nitrification of ammonia to nitrate (Daims *et al.*, 2015; Hanna *et al*, 2018).

Ammonia oxidizing microbes (AOM) obtain their energy by oxidation of ammonia (NH_3_) to nitrite (NO_2_ ^-)^. These organisms utilize a few key enzymes such as ammonia monooxygenase (AMO) and hydroxylamine oxidoreductase (HAO) to bring about the conversion. The presence of ammonia mono-oxygenase subunit-A gene (amoA) encodes ammonia mono-oxygenase (AMO), a key enzyme that catalyses the first step in ammonia oxidation. AOB was first reported in 1890 by Winogradsky and several groups began isolating and cultivating AOB from a variety of environments such as marine waters, estuarine soils and waste water treatment systems (Reddy *et al.*, 2015)

Nitrogen is an essential element for plants (Vimal *et al.*, 2017). Nitrifying bacteria play important role in soil fertility, make available nitrate nitrogen to plants (common soil nutrient element required in large quantity by plants), aid in waste treatment plant,biogeochemical cycling of nitrogen compounds and purification of the air. Nitrifying bacteria in polluted soil initiate a syntrophic pathway that provides intermediates for heterotrophic bacterial activity and thus are excellent candidates for remediation (John and Okpokwasili, 2012), With bacteria and fungi being the most important organisms for reclamation, immobilization or detoxification of metallic and radionuclide pollutants. Some bio minerals or metallic elements deposited by microbes have catalytic and other properties in nanoparticle, crystalline or colloidal forms (Gadd, 2010).

Microorganisms are very sensitive; they react quickly to any kind of changes (natural and anthropogenic) in the environment, and quickly adapt themselves to new conditions. Microorganisms take heavy metals into the cell in significant amounts. This phenomenon leads to the intracellular accumulation of metal cations of the environment and is defined as bioaccumulation (Wolejko *et al.*, 2016). Some bacterial plasmids contain specific genes for resistance to toxic heavy metal ions (Liu *et al.*, 2018; Pacwa-Plociniczak *et al.*, 2018; Lukina *et al.*, 2016; Sharma, 2016) and ability to solubilize phosphate (biofertilizers) (Ibiene *et al.*, 2012; Gupta *et al.*, 2014). Some microorganisms can adjust their metabolic activity or community structure to adapt to the harmful shock loadings. Microorganisms play important role in stress environment and the derived ecosystem functions (Singh *et al.*, 2016a, b, c; Vimal *et al.*, 2017).

Microorganisms can mobilize or immobilize metals by biosorption, sequestration, production of chelating agents, chemoorganotrophic and autotrophic leaching, methylation and redox transformations. These mechanisms stem from prior exposure of microorganisms to metals which enable them to develop the resistance and tolerance useful for biological treatment (Viti *et al.*, 2003). Microbe-metal interaction in soil/waste disposal is of interest to environmentalists in order to use adapted microorganisms as a source of biomass for bioremediation of heavy metals (Sharma, 2016; Singh *et al.*, 2016a, b, c). Metals detoxification through resistance and tolerance, this resistance can be attributed to mechanisms of exclusion or tolerance (Klassen *et al.* 2000). In an endeavour to safeguard the susceptible cellular components, a cell is capable of building up resistance to metals. Bruins *et al.* (2000) hypothesized five mechanisms for resistance to metal toxicity. These are: (1) active or dynamic transport, (2) development of a permeability barrier, (3) enzymatic detoxification, (4) reduction in sensitivity and (5) sequestration. Microbes are capable of using one or more of these methods to eliminate nonessential metals and normalize essential metals concentrations in their cells. AOB can adapt to energy stress conditions, including low nitrogen levels and low pH (Valentine, 2007). Ammonia oxidizers groups are often considered as models for unraveling the significance of microbial diversity on the responses of soil processes to environmental stress (Philippot and Hallin 2005; Wittebolle *et al.* 2009).

## MATERIALS AND METHODS

### Sample collection

Surface soil samples at depth of 0-15 cm were collected at random from Akwa Ibom State University in Akwa Ibom State, and soil sample from University of Nigeria, Nsukka. And from solid waste disposal site in Uyo, Akwa Ibom State. The soil was collected using sterile auger borer and into sterile polyethylene bag, merged to form a composite soil sample and transferred to the laboratory for analysis.

### Microbiological Analysis

#### Preparation of samples for analyses

Precisely, 5 g of the sieved soil sample was suspended in 45 ml of sterile phosphate buffer containing 139 mg of K_2_HPO_4_ and 27 mg KH_2_PO_4_ per litre (pH 7.0) and shake at 100 rpm for 2 hrs in order to liberate the organisms into the liquid medium. (Deni and Penninck, 1999; John and Okpokwasili, 2012)

#### Preparation of media

Media preparation was carried out using Winogradsky broth medium for serial dilution of soil samples and Winogradsky solid medium for the inoculation of serially diluted soil suspension.

#### Preparation of Winogradsky broth

Winogradsky broth medium phase 1 (used for the isolation of nitrifying bacteria responsible for oxidizing ammonium to nitrite) was prepared with the following composition (g/l) in sterile distilled water: (NH_4_)_2_SO_4_, 2.0; K_2_HPO_4_, 1; MgSO_4._ .7H_2_O, 0.5; NaCl, 2.0; FeSO_4_ .7H_2_O, 0.4; CaCO_3_, 0.01. Each of ten test tubes filled with 9 ml of the Winogradsky broth media 1, autoclaved at 121 °C at 15 psi for 15 minutes and allowed to cool. The test tubes was used to carry out ten-fold serial dilutions of the soil suspension (John and Okpokwasili, 2012).

#### Preparation of Winogradsky agar media

Winogradsky agar media for nitrification phases I was prepared by adding 15.0 g agar to 1000 ml of fresh broth and sterilized at 121 °C at 15 psi for 15 minutes and allowed to cool to about 45 °C before dispersing into sterile Petri dishes (John and Okpokwasili, 2012).

#### Isolation of nitrifying bacteria from soil sample

All the plates were aseptically inoculated with 0.1 ml of the appropriate dilution of the soil suspension using spread plate technique. All the inoculated Petri dishes were incubated aerobically at room temperature (28 +2°C) for 1week and examined for growth.

#### Purification of isolates

Discrete colonies that developed on Winogradsky agar media for nitrification phases 1 after 1week of incubation was aseptically sub-cultured repeatedly on corresponding freshly prepared Winogradsky agar medium. All the inoculated Petri dishes were incubated aerobically at room temperature (28 ±2°C) for 3 - 5 days. The pure isolates was transferred to Winogradsky agar slants and stored in the refrigerator for further use.

#### Identification of isolates

Pure isolates from the corresponding agar slants was characterized and identified using morphological (cell and colonial morphology, shape, motility, and gram reaction), biochemical and physiology attributes (Holt *et al.* 1994; Cheesbrough, 2006). The molecular characterization was based on 16SrDNA sequencing (Saha *et al*., 2013).

#### Physiological Characterization of the isolate

##### Nitrite determination by Griess Method (Bhaskar and Charyulu, 2005)

*Griess Ilosvay reagent will be prepared as follows*

Solution A: 0.6 g of sulphanilic acid was dissolved in 70 ml of hot distilled water, cooled, and 20 ml of concentrated HC1 was added and volume was made up to 100 ml with distilled water.

Solution B: 0.6 g of a-naphthylamine was dissolved in 10 ml of distilled water containing 1 ml of concentrated HC1 and the volume was made up to 100ml with distilled water. Solution C: 16.4 g of sodium acetate was dissolved in 70 ml of distilled water and the volume made up to 100 ml with distilled water. The three solutions (A, B and C) was stored in dark bottles and mixed in equal parts before use.

##### Ammonium oxidation test for determination of nitrite

Five millilitres of Winogradsky mineral basal medium was prepared. The tubes was sterilized by autoclaving at 121°C at 15 psi for 15 minutes and allowed to cool. One loopful of each ammonium oxidizing bacteria isolate was added into each tube and incubated aerobically for 5 days at room temperature. At the end of the incubation period, the presence of nitrite was tested using Griess Ilosvay reagent. The reagent was added and observed for the development of purplish red/pink colouration within 5 minutes (Joel and Amajuoyi, 2010; Hoang *et al.*, 2016).

##### Inoculum preparation and standardization

Inocula were prepared by inoculating isolates onto prepared nutrient agar plates and incubating at 30°C for 24 h. After incubation, colonies were suspended in test tubes containing sterile normal saline solution. The tubes were vortex for 2 min, and then transferred into a sterile test tube. The cells suspension was adjusted to a 0.5 McFarland standard (Optical density of 0.5 - 1.0 at 600nm) using sterile normal saline to give a concentration of 10^8^ cfu/ml, to get the final inocula.

##### Tolerance study

Mineral salts medium of the following composition (g/l): (NH_4_)_2_SO_4_, 1.0; KH_2_PO_4_, 1.0 g; K_2_HPO_4_, 1.0 g; MgSO_4_, 0.2; CaCl_2_, 0.02; FeCl_3_.6H_2_O; 0.004 for ammonium oxidizing bacteria. Sterilized by autoclaving at 121°C at 15 psi for 15 minutes and allowed to cool.

Salts of Cupper (Cu), Nickel (Ni), Cadmium (Cd), and Lead (Pb) was used as CuSO_4_.5H_2_O, NiSO_4_.6H_2_O, CdCl_2._H_2_O and Pb(CH_2_COO)_2_)_2_. (OH) _2_, respectively.

##### Primary Screening of heavy metal resistant nitrifying bacteria

An amount (0.1 ml) of the standard inoculum was plated into mineral salts agar medium supplement with metal salt concentrations of 100 mg/L. The plates were incubated at room temperature for 3 - 5 days (Jamaluddin *et al.*, 2012; Pandit *et al.*, 2013). Ammonia oxidizing bacterial isolates will be selected for tolerance study.

##### Experimental Set up

Analytical grades of metal salts was used to prepare stock solutions. The mineral salt medium for ammonia oxidizing bacteria was amended with the appropriate aliquot of the stock solution of the metal salt concentrations of 100 mg/L, 200 mg/L, 500 mg/L and 1000 mg/L.

##### Effect of heavy metal on the growth of nitrifying isolates

Changes in population of the nitrifying isolates were monitored following their exposure to heavy metals. About 1 ml of the standard inoculum was introduced into each flask containing heavy metal salts amended in mineral salt medium (MSM). The cultures were incubated aerobically at room temperature (28 ±2°C) for 7 days. The growth was measured by withdrawing samples from the medium every 24 hours and absorbance of the turbidity measured at 600 nanometre using spectrophotometer (Bhastar and Charyulu, 2005).

##### Statistical Analysis

Result reported as mean ± standard deviation. All data were subjected to statistical analysis by analysis of variance (ANOVA). The means were separated with least significant difference. The result consider significant at P < 0.05. Least significant difference test (LSD) was also being performed between each treatment and the control. Correlation (association) and regression (changes) analysis was done using statistical product and service solution (SPSS) for windows version 20.

## RESULTS

### Morphological and Biochemical Characterization of Ammonia Oxidizing Bacterial Isolates

Table 1 shows the morphological and biological characteristic of ammonia oxidizing isolate, the five potential heavy metal tolerance Ammonia oxidizing bacteria were characterized based on their cultural, morphological and biochemical characteristic. Isolate were identified as Gram negative; Catalase positive; Indole negative; Methylene red negative; Voges proskauer negative; Urease negative and Oxidase negative. The isolates were compared with Standard description of Bergey’s Manual of determinative bacteriology. A1, A2, A3, A4 and A5 represent different Ammonia oxidizing bacteria.

**Table 1:** Morphological and Biochemical Characterization of Ammonia Oxidizing Bacterial Isolates.

| Suspected organism | Colony | Cell shape | Gram stain | Spore stain | Cat | H<sub>2</sub>S | Ind | Cit | MR | Vp | Ur | Mot | Oxid | Nit red |
| --- | --- | --- | --- | --- | --- | --- | --- | --- | --- | --- | --- | --- | --- | --- |
| A1 | Colourless mucoid raise, | Rod | - | - | + | - | - | + | - | - | + | + | + | + |
| A2 | White mucoid, raise | Rod | - | - | + | - | - | + | - | + | - | + | + | + |
| A3 | White Mucoid raise, | Rod | - | - | + | - | - | + | - | - | - | + | + | + |
| A4 | Raise,<br>brownish | Rod | - | - | + | + | - | + | - | + | - | - | - | + |
| A5 | Colourless<br>mucoid<br>raise, | Rod | - | - | + | + | + | - | + | - | + | + | + | + |

+ present (positive)
- absent (negative)
Cat Catalase
Ind Indole
Cit Citrate
MR Methyl red
Vp Voges proskauer
Ur Urease
Mot Motility

### Tolerance of isolates to the copper

The results represented in Figures 1, 2 3 and 4 shows the level of tolerance at the different metal salt concentrations by the ammonia oxidizing bacteria isolates. Figure 1 shows the tolerance of the A1 isolates for different concentrations of Cu. The growth of A1 range from 0.0535 ± .00071 to 0.0725±.00071 within 72 hours, increase to 0.0935±0.0072 within 120hrs and decrease to 0.0715 ± 0.00212 after 168 hours at 0 concentration. .. A1 growth range from 0.0515 ± .000354 to 0.043±.00424 within 72 hours, increase to 0.0735±0.00212 within 120hrs and decrease to 0.05 ± 0.00283 after 168 hours at 100ug/ml concentration. 0.0365 ± .00071 to 0.036±.0001 within 72 hours; decrease to 0.0205±0.001 within 120hrs and decrease to 0.006 ± 0.00283 after 168 hours at 1000ug/ml concentration.

Figure 2 shows the tolerance of the A2 isolates for different concentrations of Cu. The growth of A2 range from 0.057 ± .000141 to 0.086±.00283 within 72 hours, increase to 0.1115±0.00276 within 120hrs and decrease to 0.078 ± 0.00424 after 168 hours at 0 concentration. A2 growth range from 0.0535 ± .0078 to 0.0525±.00212 within 72 hours, increase to 0.0865±0.00354 within 120hrs and decrease to 0.058 ± 0.018 after 168 hours at 100ug/ml concentration. 0.032 ± .00141 to 0.0275±.00212 within 72 hours; decrease to 0.026±0.0028 within 120hrs and decrease to 0.095 ± 0.00212 after 168 hours at 100ug/ml concentration.

Figure 3 shows the tolerance of the A3 isolates for different concentrations of Cu. The growth of A3 range from 0.0685 ± .0001 to 0.0755±.0.001 within 72 hours, increase to 0.0865±0.001 within 120hrs and decrease to 0.083 ± 0.008 after 168 hours at 0 concentration. A3 growth range from 0.0645 ± .004 to 0.0725±.001 within 72 hours; increase to 0.0745±0.001 within 120hrs and decrease to 0.0535 ± 0.006 after 168 hours at 100ug/ml concentration. A3 growth range from 0.0295 ± .001 to 0.0215±.001 within 72 hours, increase to 0.0215 ±0.003 within 120hrs and decrease to 0.015 ± 0.003 after 168 hours at 1000ug/ml concentration.

Figure 4 shows the tolerance of the A4 isolates for different concentrations of Cu. The growth of A4 range from 0.0655 ± .002 to 0.0835 ±.002 within 72 hours, increase to 0.095 ±0.001 within 120hrs and decrease to 0.072 ± 0.013 after 168 hours at 0 concentration. . A4 growth range from 0.0635 ± .002 to 0.064 ±.004 within 72 hours; increase to 0.081±0.001 within 120hrs and decrease to 0.0445 ± 0.001 after 168 hours at 100ug/ml concentration. A4 growth range from 0.032 ± .00 to 0.023 ±.000 within 72 hours, increase to 0.023 ±0.004 within 120hrs and decrease to 0.016 ± 0.001 after 168 hours at 1000ug/ml concentration.

Figure 5 shows the tolerance of the A5 isolates for different concentrations of Cu. The growth of A5 range from 0.062 ± .000141 to 0.0795 ±.001 within 72 hours, increase to 0.084 ± 0.001 within 120hrs and decrease to 0.0645 ± 0.008 after 168 hours at 0 concentration. A5 growth range from 0.0585 ± .005 to 0.068±.007 within 72 hours, increase to 0.0795 ± 0.007 within 120hrs and decrease to 0.0545 ± 0.008 after 168 hours at 100ug/ml concentration. A5 growth range from 0.03 ± .008 to 0.026 ±.000 within 72 hours; increase to 0.026±0.0028 within 120hrs and decrease to 0.018 ± 0.003 after 168 hours at 1000ug/ml concentration.

The results represented in Figures 6, 7, 8, 9 and 10 shows the level Response of different Ammonia Oxidizing Bacteria on different concentration of Copper within 24, 72, 120 and 168 hour respectively.

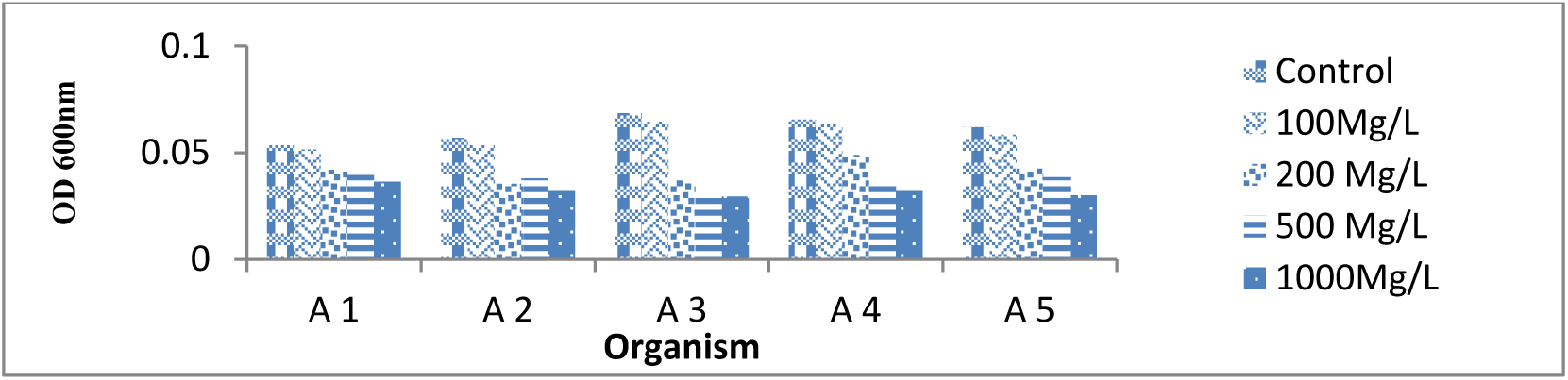
Response of different Ammonia Oxidizing Bacteria on different concentration of Copper within 24 hours.

**Figure 7:**
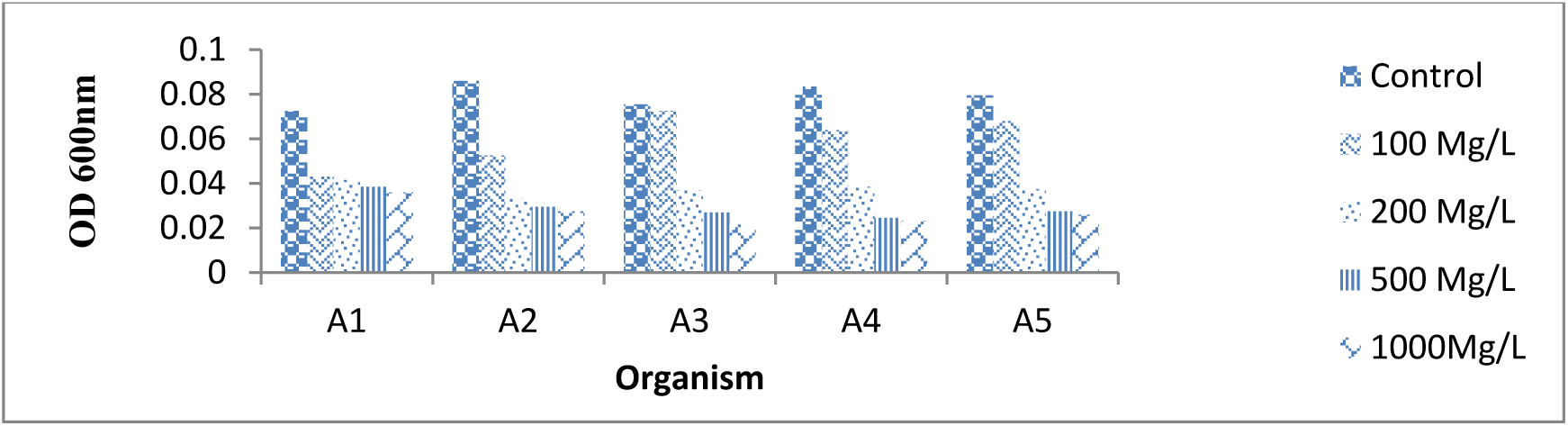
Response of different Ammonia Oxidizing Bacteria on different concentration of Copper within 72 hours.

**Figure 8:**
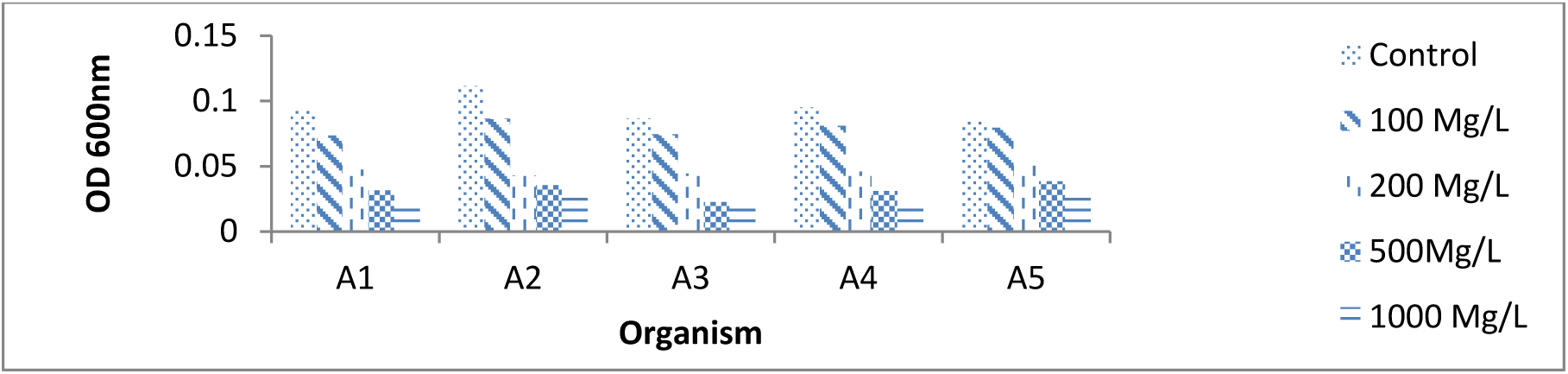
Response of different Ammonia Oxidizing Bacteria on different concentration of Copper within 120 hrs.

**Figure 9:**
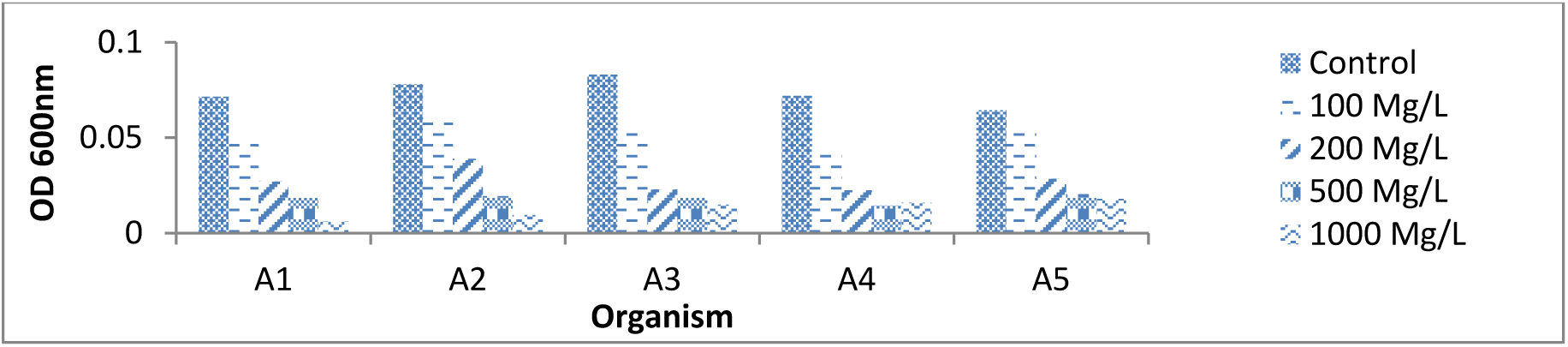
Response of different Ammonia Oxidizing Bacteria on different concentration of Copper within 168 hours.

**Figure 10:**
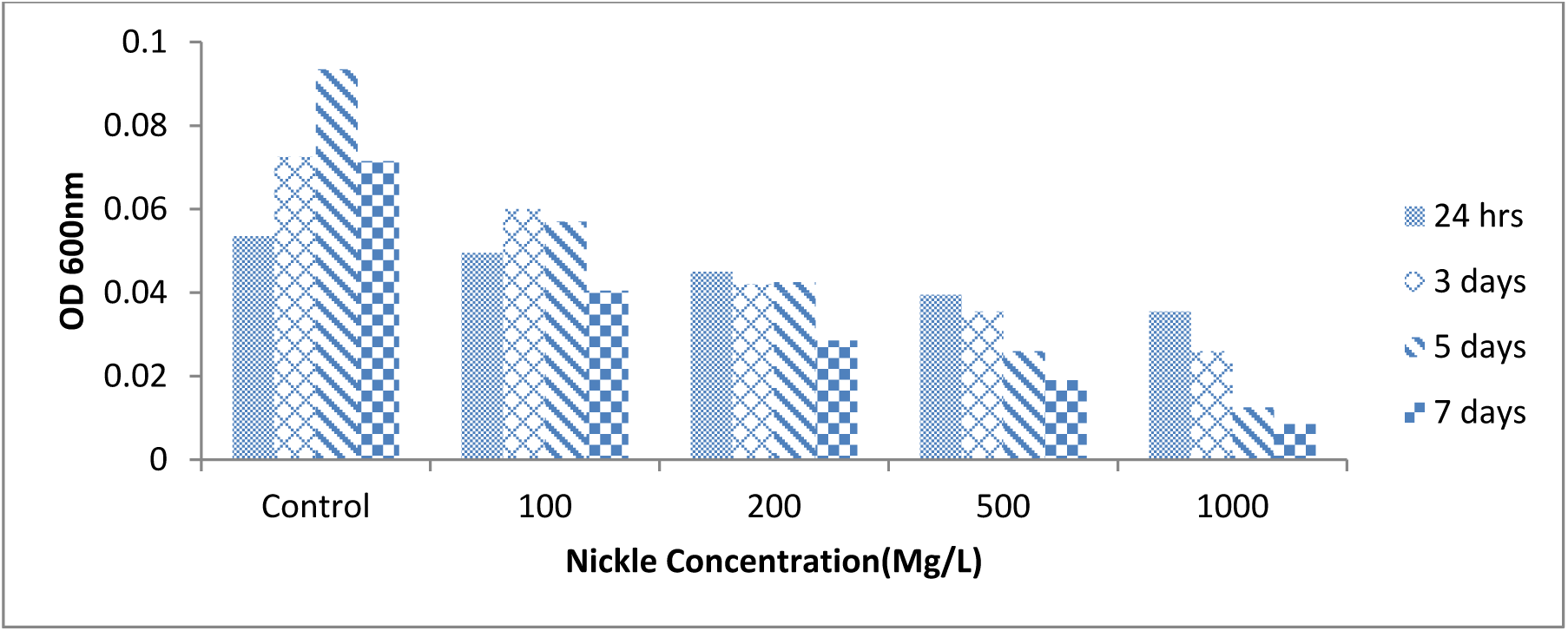
Tolerance of A1 to different nickel concentration.

### Tolerance of isolates to the Nickel

The results represented in Figures 10, 11, 12, 13 and 14 shows the level of tolerance at the different nickel concentrations by the ammonia oxidizing bacteria isolates.

**Figure 11:**
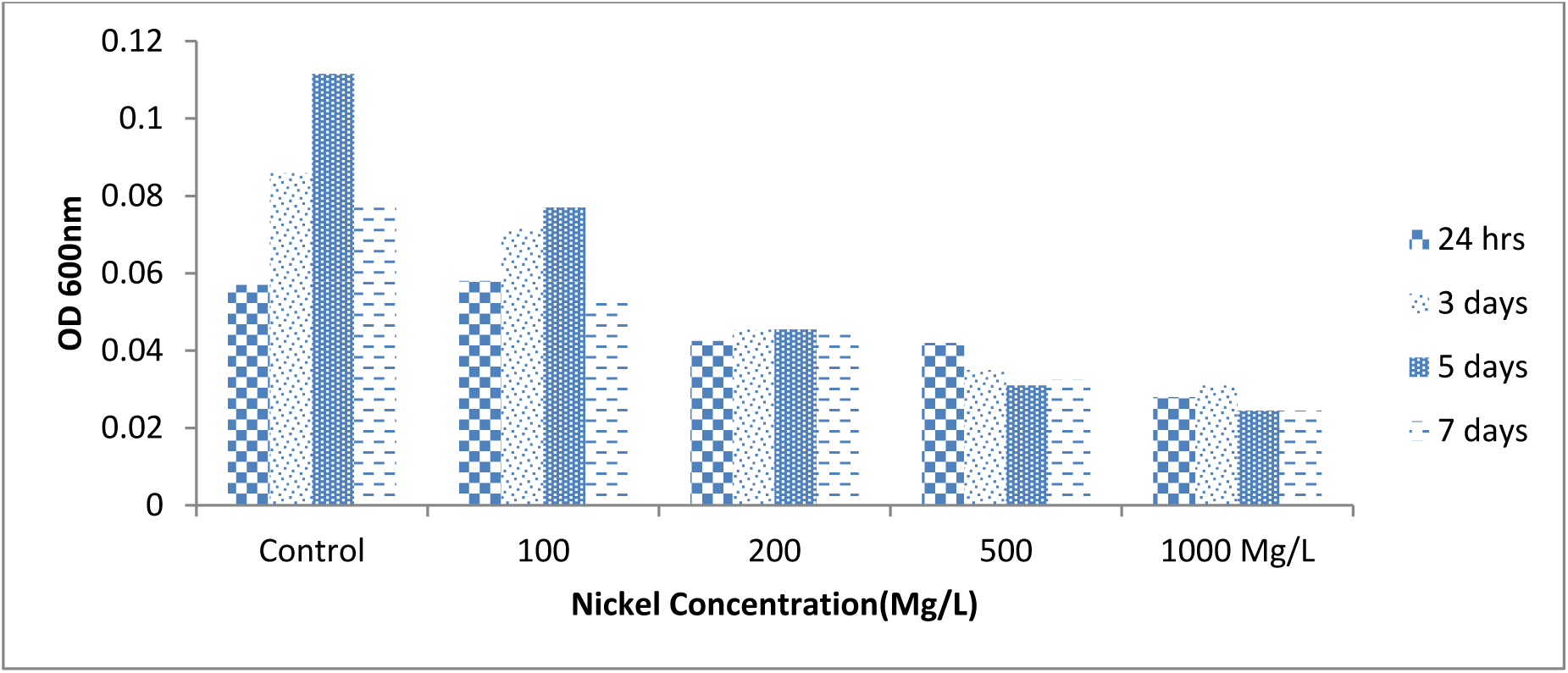
Tolerance of A2 to different nickel concentration.

**Figure 12:**
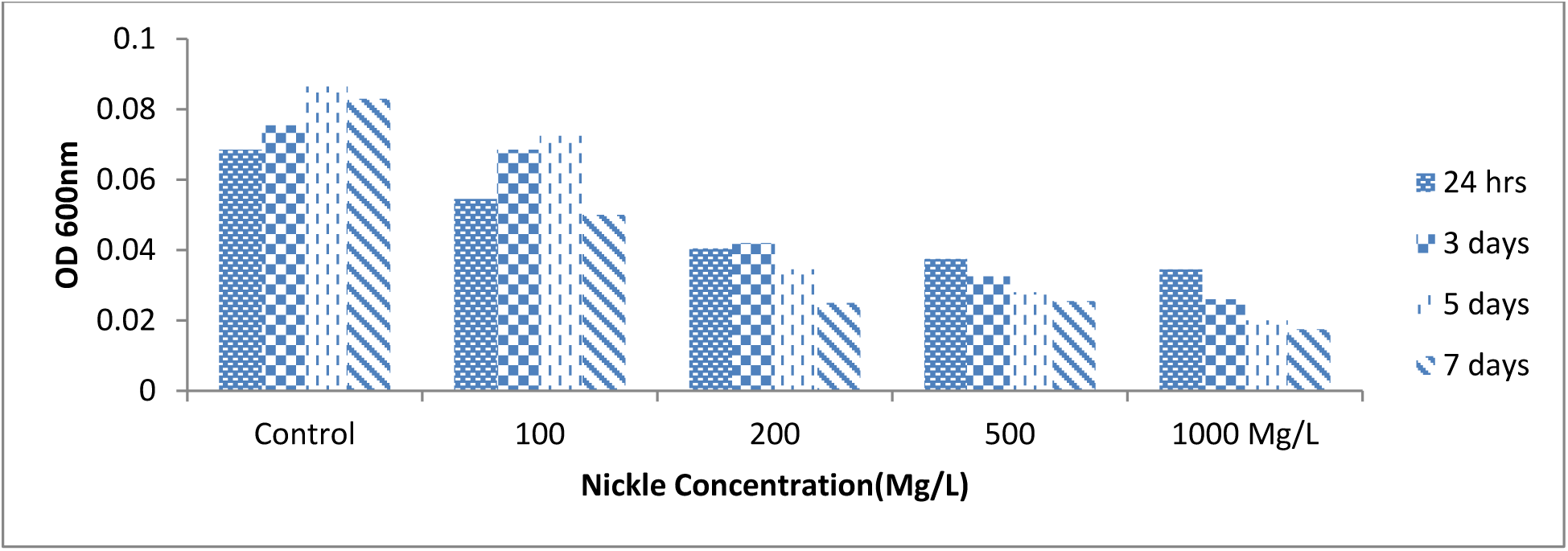
Tolerance of A3 to different nickel concentration.

**Figure 13:**
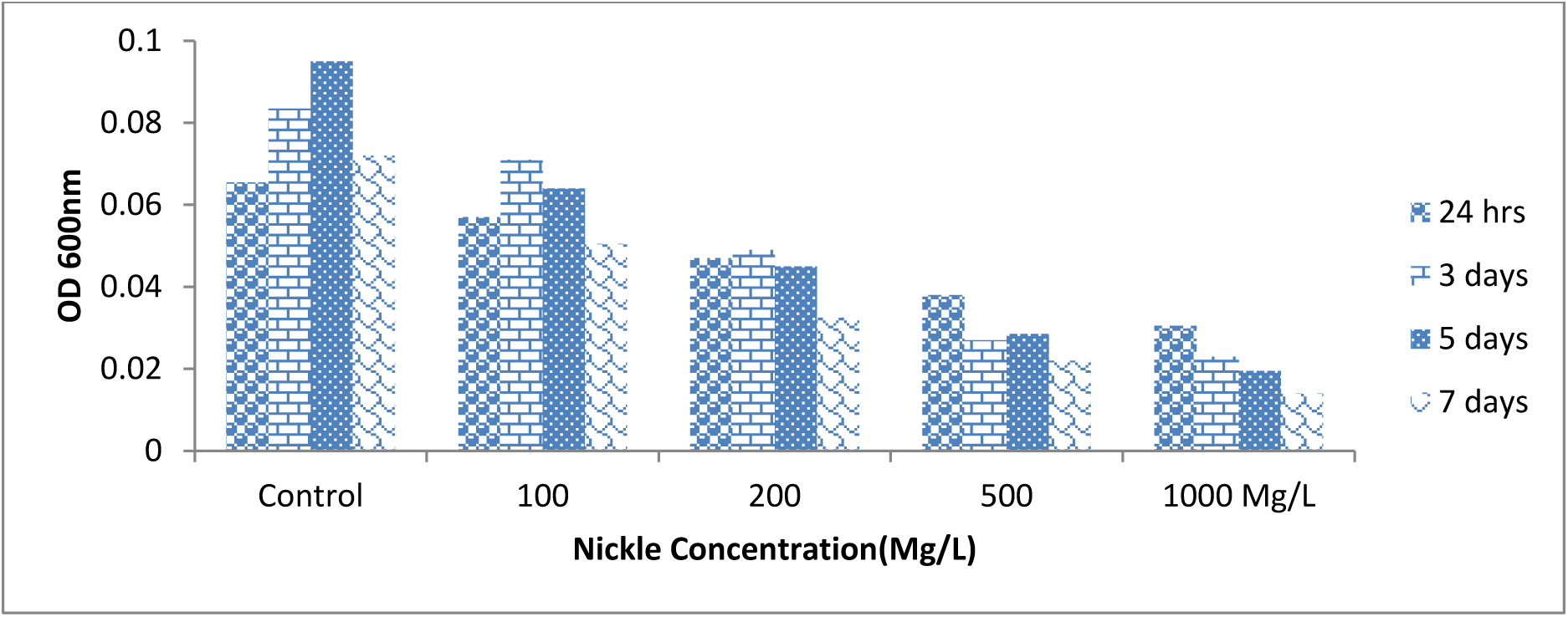
Tolerance of A4 to different nickel concentration.

**Figure 14:**
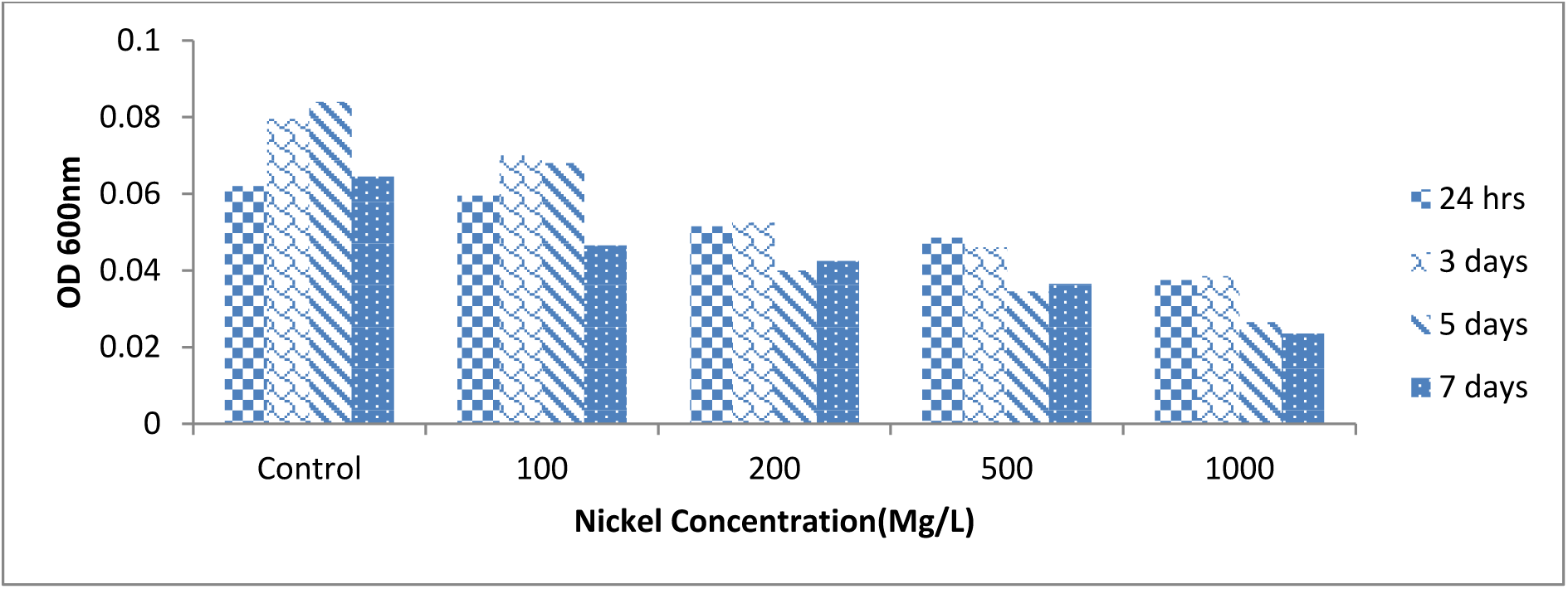
Tolerance of A5 to different nickel concentration.

Figure 10 shows the tolerance of the A1 isolates for different concentrations of Ni. The growth of A1 range from 0.0535 ± .00071 to 0.0725±.00071 within 72 hours, increase to 0.0935±0.0072 within 120hrs and decrease to 0.0715 ± 0.00212 after 168 hours at 0 concentration. A1 growth range from 0.0495 ± .0001 to 0.06±.0.00 within 72 hours, decrease to 0.057 ±0.008 within 120hrs and decrease to 0.0405 ± 0.002 after 168 hours at 100ug/ml concentration. 0.0355 ± .006 to 0.026±.0.00 within 72 hours; decrease to 0.0125 ±0.002 within 120hrs and decrease to 0.0085 ± 0.001 after 168 hours at 1000ug/ml concentration.

Figure 11 shows the tolerance of the A2 isolates for different concentrations of Ni. The growth of A2 range from 0.057 ± .000141 to 0.086±.00283 within 72 hours, increase to 0.1115±0.00276 within 120hrs and decrease to 0.078 ± 0.00424 after 168 hours at 0 concentration. A2 growth range from 0.058 ± .0004 to 0.0715 ±.0.01 within 72 hours, increase to 0.077 ± 0.014 within 120hrs and decrease to 0.078 ± 0.004 after 168 hours at 100ug/ml concentration. A2 growth range from 0.028 ± 0.004 to 0.031± 0.004 within 72 hours; decrease to 0.0245 ± 0.018 within 120hrs; 0.0245 ± 0.006 after 168 hours at 1000ug/ml concentration.

Figure 12 shows the tolerance of the A3 isolates for different concentrations of Ni. The growth of A3 range from 0.0685 ± .0001 to 0.0755±.0.001 within 72 hours, increase to 0.0865±0.001 within 120hrs and decrease to 0.083 ± 0.008 after 168 hours at 0 concentration. A3 growth range from 0.0545 ± .005 to 0.0685 ±.002 within 72 hours; Increase to 0.0725 ± 0.001 within 120hrs and increase to 0.083 ± 0.008 after 168 hours at 100ug/ml concentration. A3 growth range from 0.0345 ± .0026 to 0.026 ± 0.00 within 72 hours, decrease to 0.02 ±0.013 within 120hrs and decrease to 0.0175 ± 0.004 after 168 hours at 1000ug/ml concentration.

Figure 13 shows the tolerance of the A4 isolates for different concentrations of Ni. The growth of A4 range from 0.0655 ± .002 to 0.0835 ±.002 within 72 hours, increase to 0.095 ±0.001 within 120hrs and decrease to 0.072 ± 0.013 after 168 hours at 0 concentration. . A4 growth range from 0.057 ± 0.003 to 0.071 ± 0.01 within 72 hours; increase to 0.064 ± 0.004 within 120hrs and increase to 0.072 ± 0.011 after 168 hours at 100ug/ml concentration. A4 growth range from 0.0305 ± 0.001 to 0.023 ± 0.007 within 72 hours, decrease to 0.0195 ±0.006 within 120hrs and decrease to 0.014 ± 0.004 after 168 hours at 1000ug/ml concentration.

Figure 14 shows the tolerance of the A5 isolates for different concentrations of Ni. The growth of A5 range from 0.062 ± .000141 to 0.0795 ±.001 within 72 hours, increase to 0.084 ± 0.001 within 120hrs and decrease to 0.0645 ± 0.008 after 168 hours at 0 concentration. A5 growth range from 0.0595 ± 0.005 to 0.07 ± 0.001 within 72 hours, decrease to 0.068 ± 0.007 within 120hrs and decrease to 0.0645 ± 0.007 after 168 hours at 100ug/ml concentration. A5 growth range from 0.0375 ± .004 to 0.0385 ± 0.016 within 72 hours; decrease to 0.0265 ±0.002 within 120hrs and decrease to 0.0235 ± 0.001 after 168 hours at 1000ug/ml concentration.

The results represented in Figures 15, 16, 17 and 18 shows the level Response of different Ammonia Oxidizing Bacteria on different concentration of Nickel within 24, 72, 120 and 168 hour respectively.

**Figure 15:**
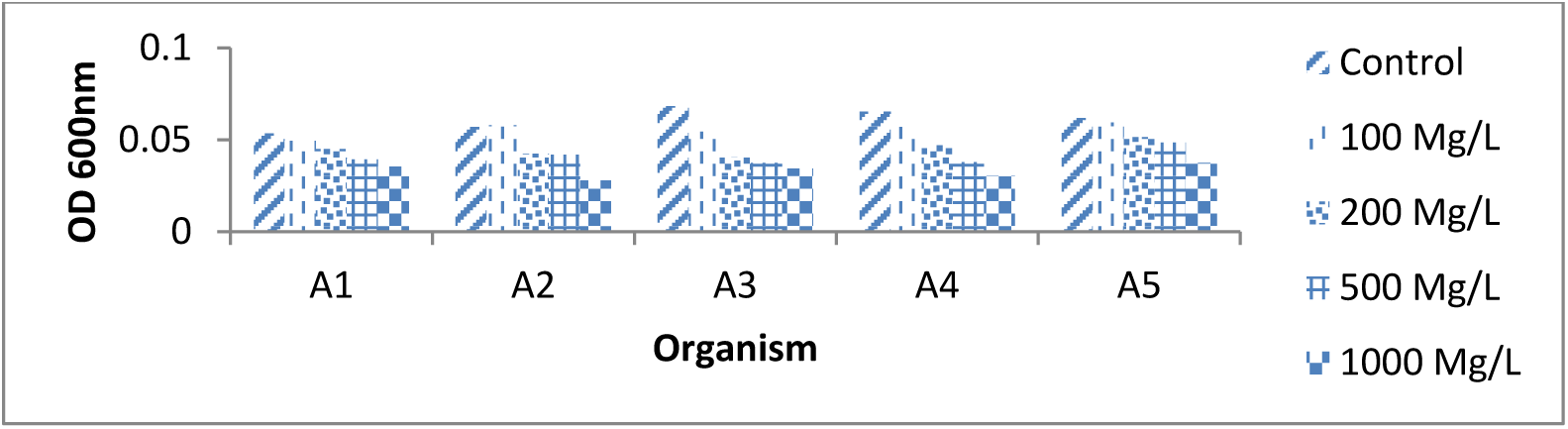
Response of different Ammonia Oxidizing Bacteria on different concentration of Nickel within 24 hours.

**Figure 16:**
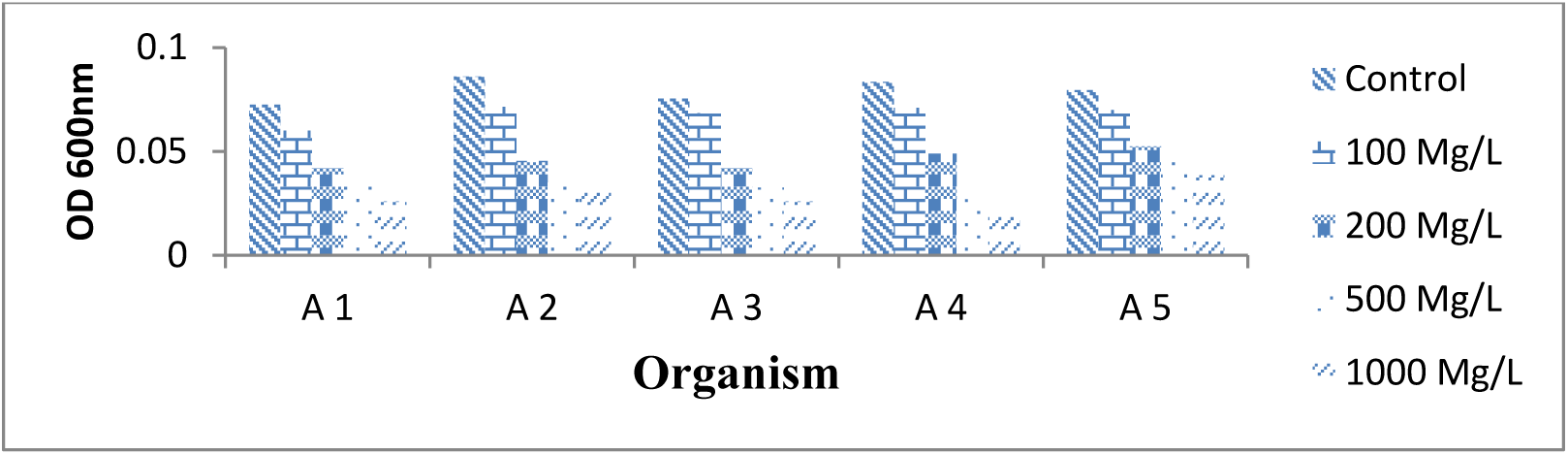
Response of different Ammonia Oxidizing Bacteria on different concentration of Nickel within 72 hours.

**Figure 17:**
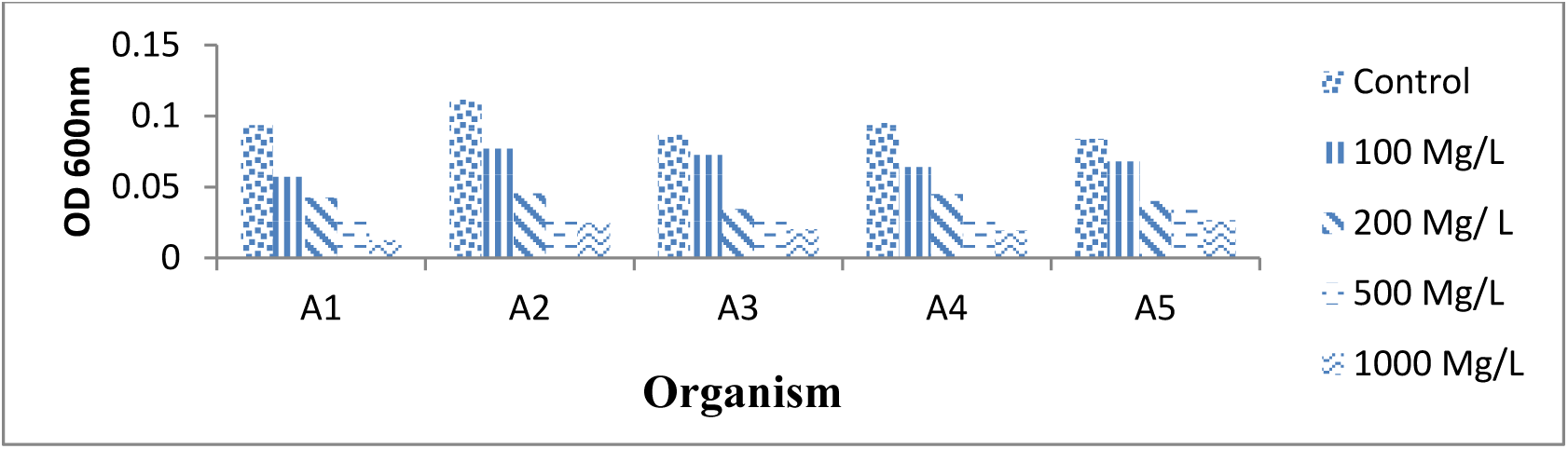
Response of different Ammonia Oxidizing Bacteria on different concentration of Nickel within 120 hours.

**Figure 18:**
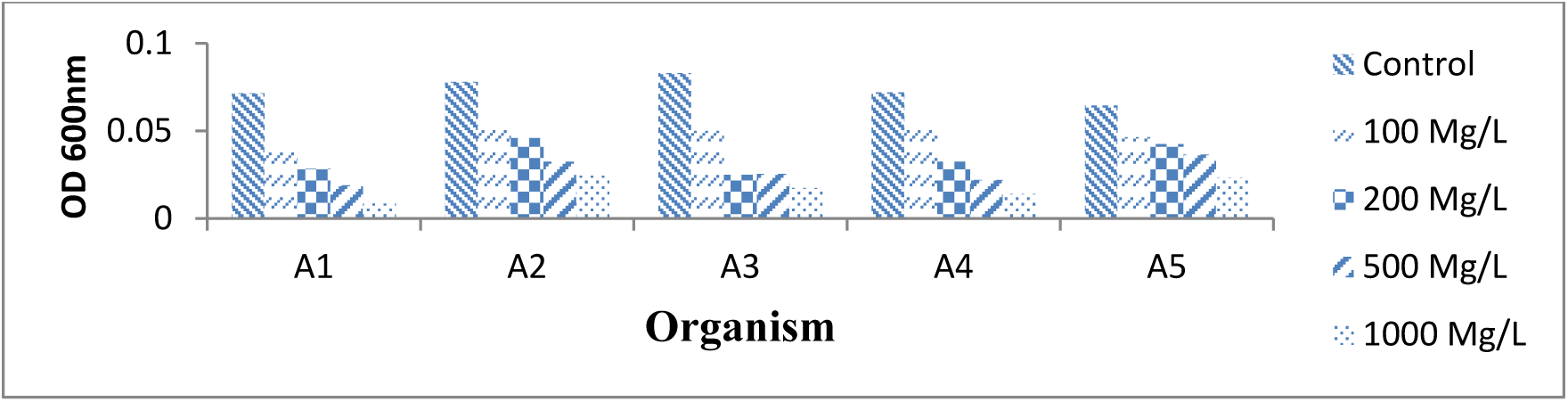
Response of different Ammonia Oxidizing Bacteria on different concentration of Nickel within168 hours.

### Tolerance of isolates to the lead

The results represented in Figures 19, 20 21, 22 and 23 shows the level of tolerance at the different lead concentrations by the ammonia oxidizing bacteria isolates.

**Figure 19:**
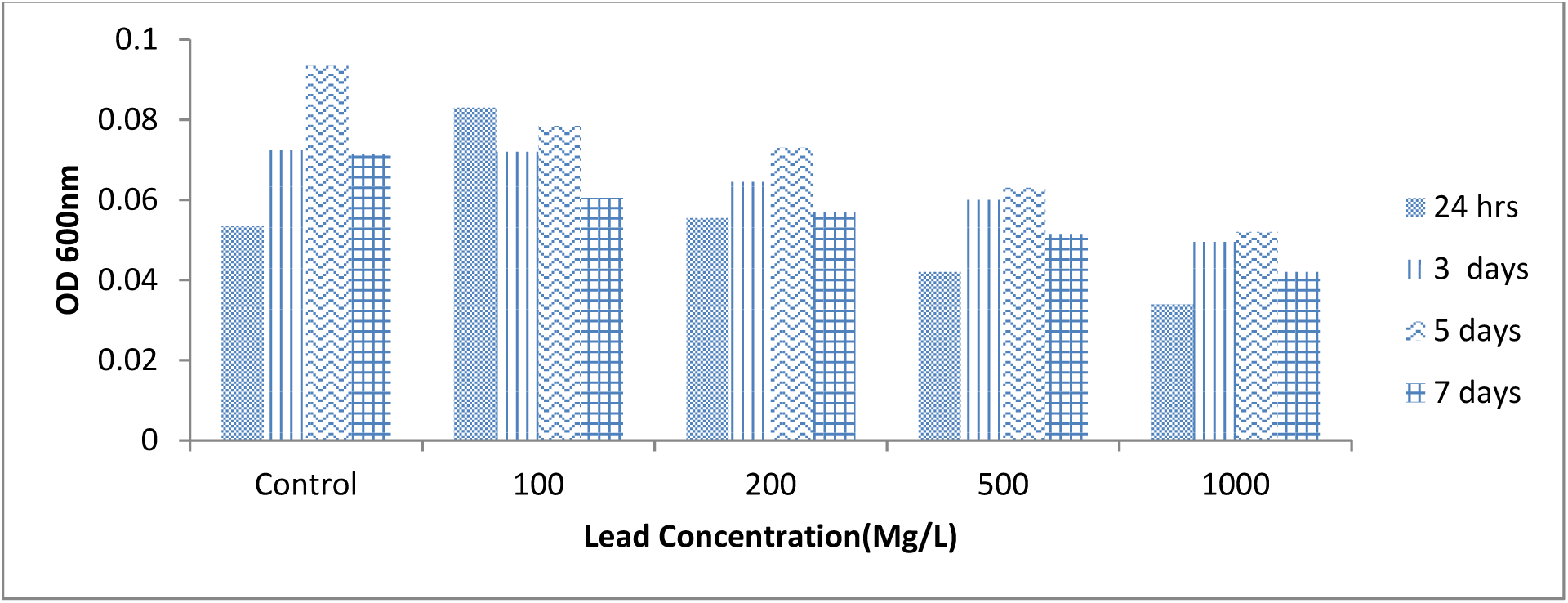
Tolerance of A1 to different lead concentration.

**Figure 20:**
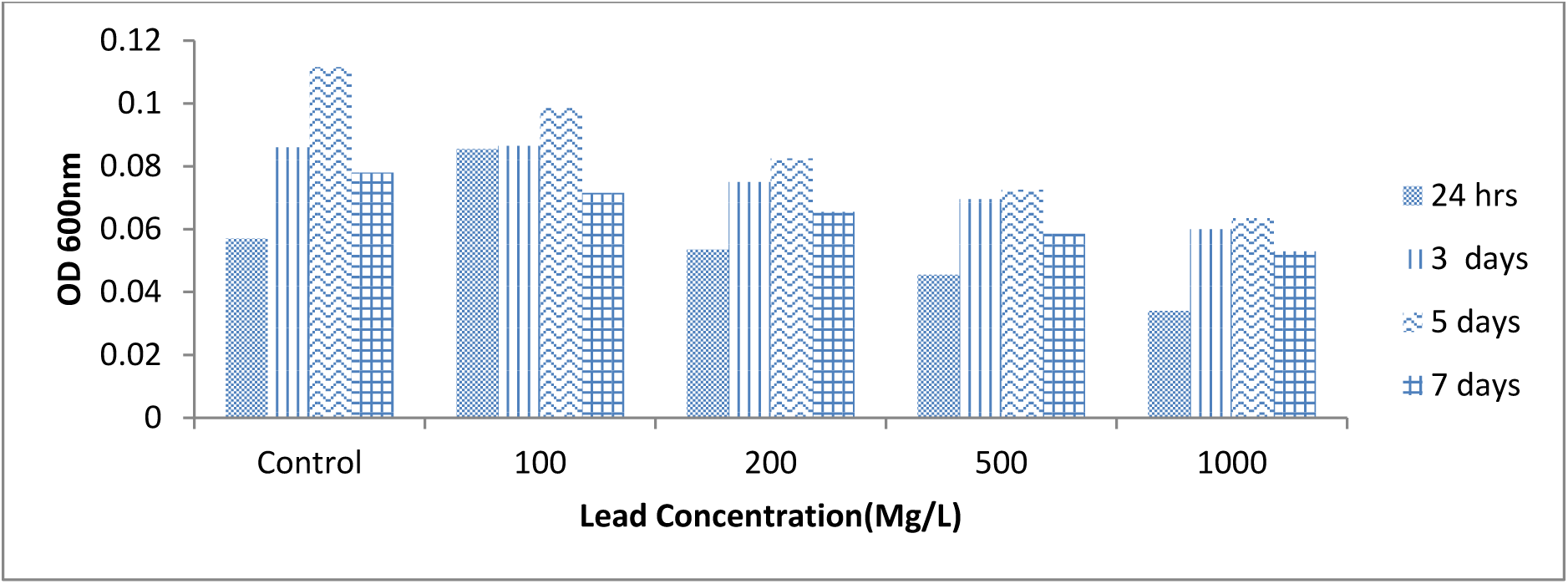
Tolerance of A2 to different lead concentration.

**Figure 21:**
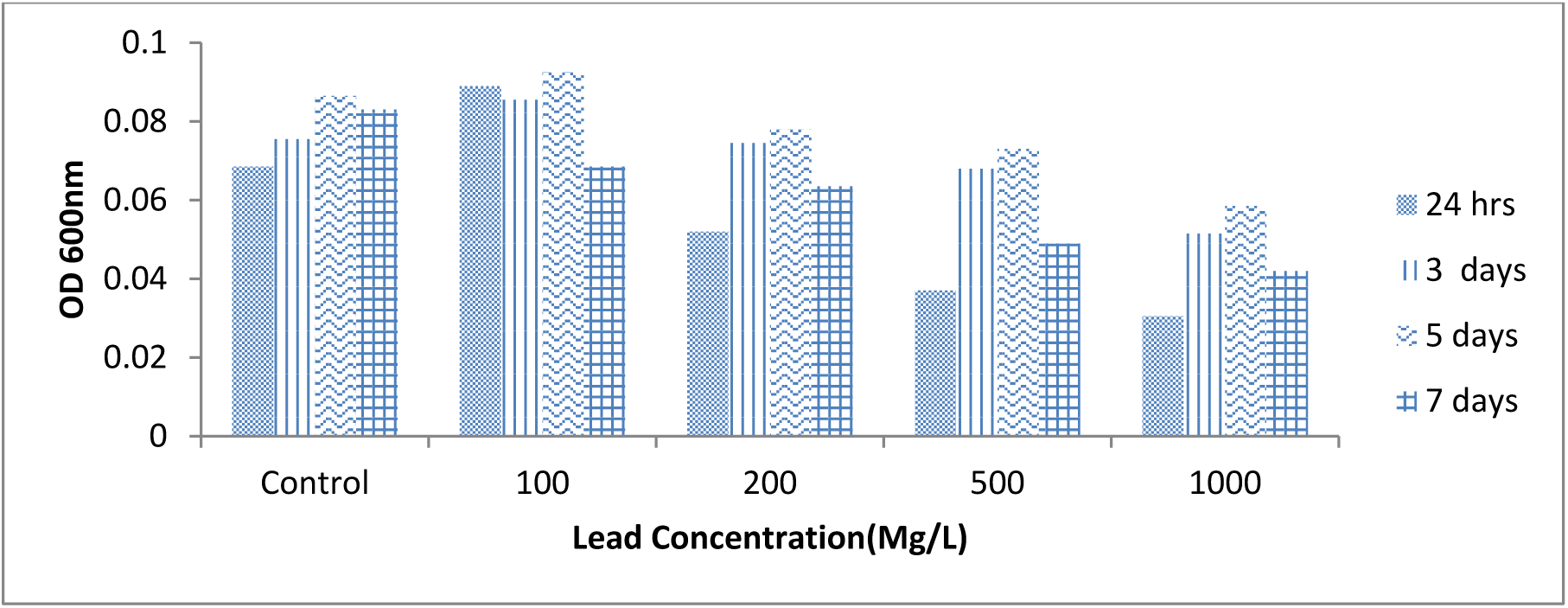
Tolerance of A3 to different lead concentration.

**Figure 22:**
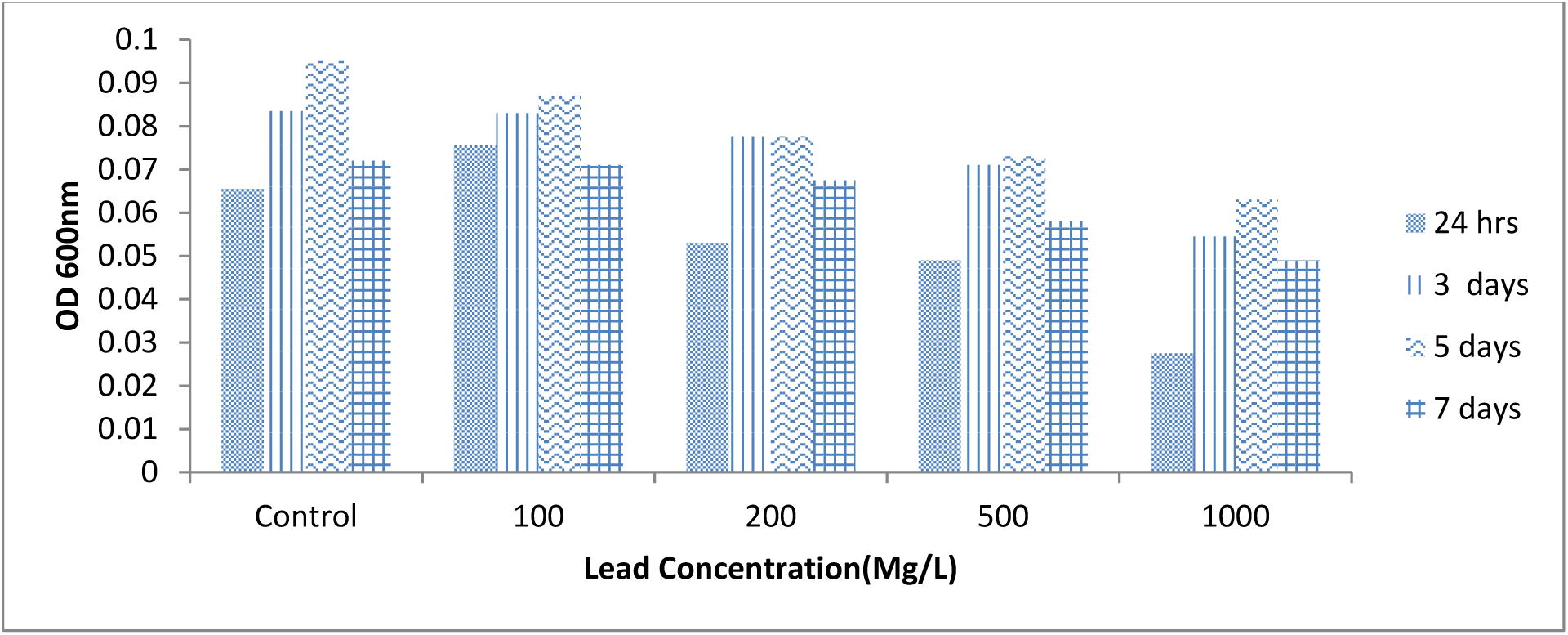
Tolerance of A4 to different lead concentration.

**Figure 23:**
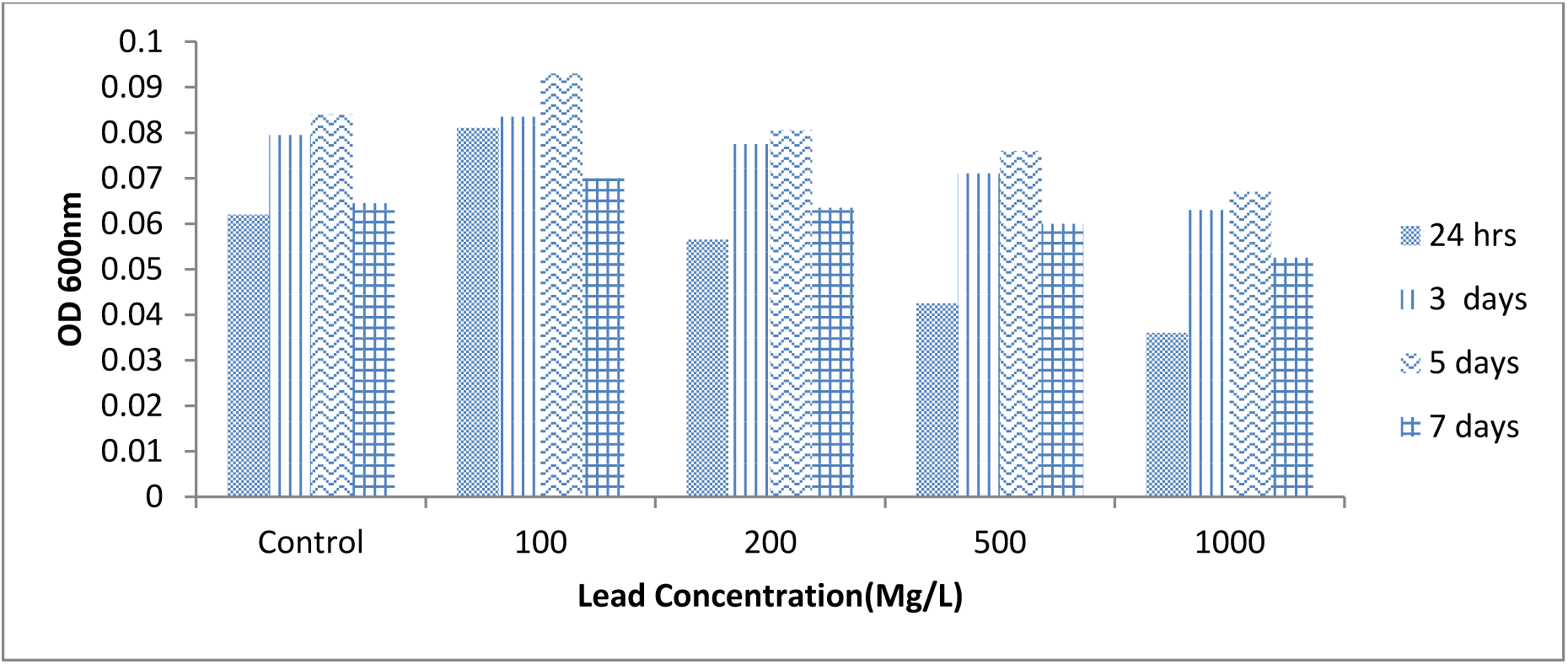
Tolerance of A5 to different lead concentration.

Figure 19 shows the tolerance of the A1 isolates for different concentrations of Pb. The growth of A1 range from 0.0535 ± .00071 to 0.0725±.00071 within 72 hours, increase to 0.0935±0.0072 within 120hrs and decrease to 0.0715 ± 0.00212 after 168 hours at 0 concentration. A1 growth range from 0.083 ± 0.007 to 0.072 ±.0.001 within 72 hours, increase to 0.0785 ±0.001 within 120hrs and decrease to 0.0605 ± 0.001 after 168 hours at 100ug/ml concentration. A1 growth range from 0.034 ± .001 to 0.0495±.0.001 within 72 hours; increase to 0.052 ±0.001 within 120hrs and decrease to 0.0042 ± 0.003 after 168 hours at 1000ug/ml concentration.

Figure 20 shows the tolerance of the A2 isolates for different concentrations of Pb. The growth of A2 range from 0.057 ± .000141 to 0.086±.00283 within 72 hours, increase to 0.1115±0.00276 within 120hrs and decrease to 0.078 ± 0.00424 after 168 hours at 0 concentration. A2 growth range from 0.0855 ± 0.001 to 0.0985 ±.0.001 within 72 hours, decrease to 0.0715 ± 0.001 within 120hrs and decrease to 0.089 ± 0.00 after 168 hours at 100ug/ml concentration. A2 growth range from 0.034 ± 0.008 to 0.06± 0.004 within 72 hours; increase to 0.0635 ± 0.005 within 120hrs; 0.053 ± 0.008 after 168 hours at 1000ug/ml concentration.

Figure 21 shows the tolerance of the A3 isolates for different concentrations of Pb. The growth of A3 range from 0.0685 ± .0001 to 0.0755±.0.001 within 72 hours, increase to 0.0865±0.001 within 120hrs and decrease to 0.083 ± 0.008 after 168 hours at 0 concentration. A3 growth range from 0.089 ± 0.00 to 0.0855 ± 0.006 within 72 hours; Increase to 0.0925 ± 0.009 within 120hrs and decrease to 0.0685 ± 0.002 after 168 hours at 100ug/ml concentration. A3 growth range from 0.0305 ± 0.002 to 0.0515 ± 0.01 within 72 hours, decrease to 0.0585 ± 0.016 within 120hrs and decrease to 0.042 ± 0.018 after 168 hours at 1000ug/ml concentration.

Figure 22 shows the tolerance of the A4 isolates for different concentrations of Pb. The growth of A4 range from 0.0655 ± .002 to 0.0835 ±.002 within 72 hours, increase to 0.095 ±0.001 within 120hrs and decrease to 0.072 ± 0.013 after 168 hours at 0 concentration. . A4 growth range from 0.0755 ± 0.006 to 0.083 ± 0.003 within 72 hours; increase to 0.087 ± 0.004 within 120hrs and decrease to 0.071 ± 0.01 after 168 hours at 100ug/ml concentration. A4 growth range from 0.0275 ± 0.019 to 0.0545 ± 0.008 within 72 hours, increase to 0.063 ± 0.001 within 120hrs and decrease to 0.049 ± 0.001 after 168 hours at 1000ug/ml concentration.

Figure 23 shows the tolerance of the A5 isolates for different concentrations of Pb. The growth of A5 range from 0.062 ± .000141 to 0.0795 ±.001 within 72 hours, increase to 0.084 ± 0.001 within 120hrs and decrease to 0.0645 ± 0.008 after 168 hours at 0 concentration. A5 growth range from 0.081 ± 0.001 to 0.0835 ± 0.004 within 72 hours, increase to 0.093 ± 0.004 within 120hrs and decrease to 0.07 ± 0.001 after 168 hours at 100ug/ml concentration. A5 growth range from 0.036 ± 0.008 to 0.063 ± 0.002 within 72 hours; decrease to 0.067 ±0.001 within 120hrs and decrease to 0.0525 ± 0.005 after 168 hours at 1000ug/ml concentration.

The results represented in Figures 24, 25, 25, 26 and 27 shows the level Response of different Ammonia Oxidizing Bacteria on different concentration of lead within 24, 72, 120 and 168 hour respectively.

**Figure 24:**
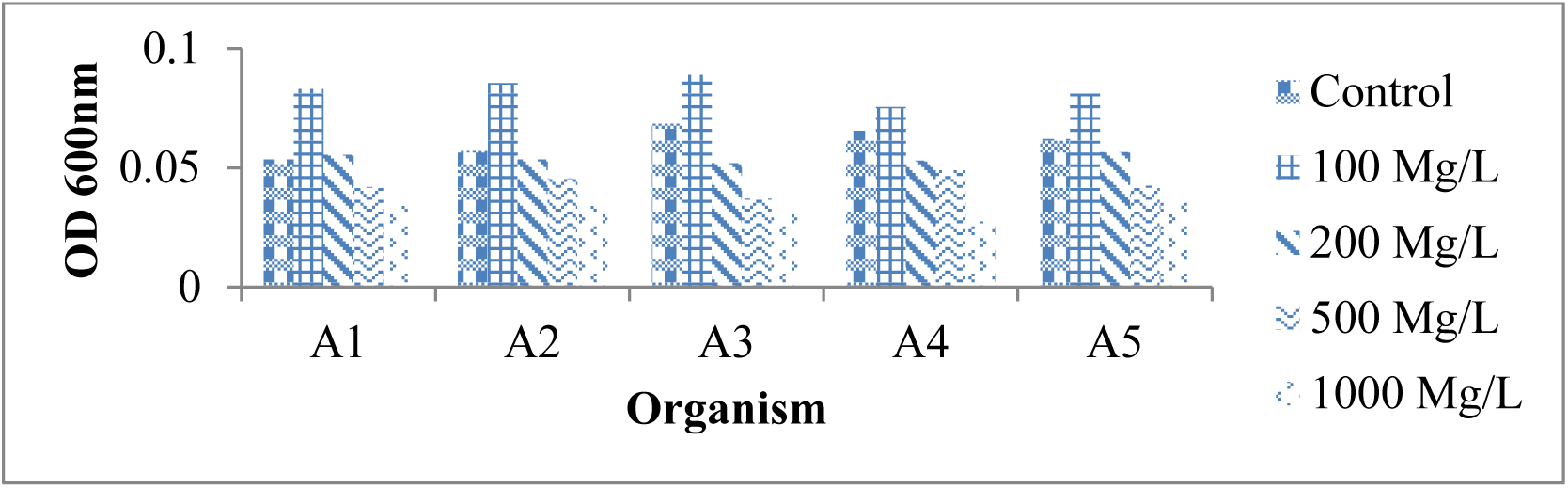
Response of different Ammonia Oxidizing Bacteria on different concentration of Lead within 24 hours.

**Figure 25:**
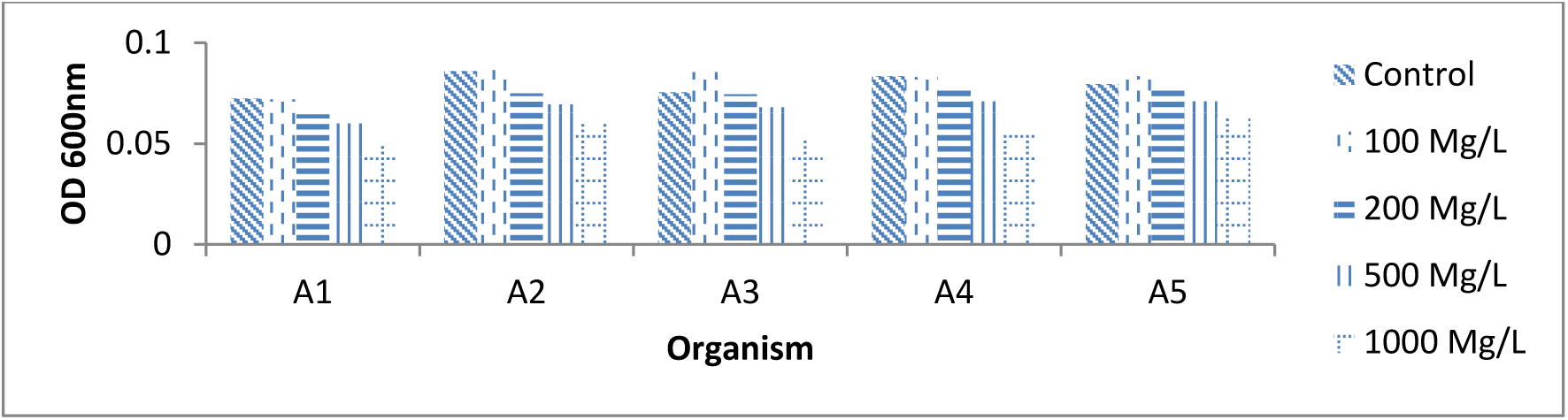
Response of different Ammonia Oxidizing Bacteria on different concentration of Lead within 72 hours.

**Figure 26:**
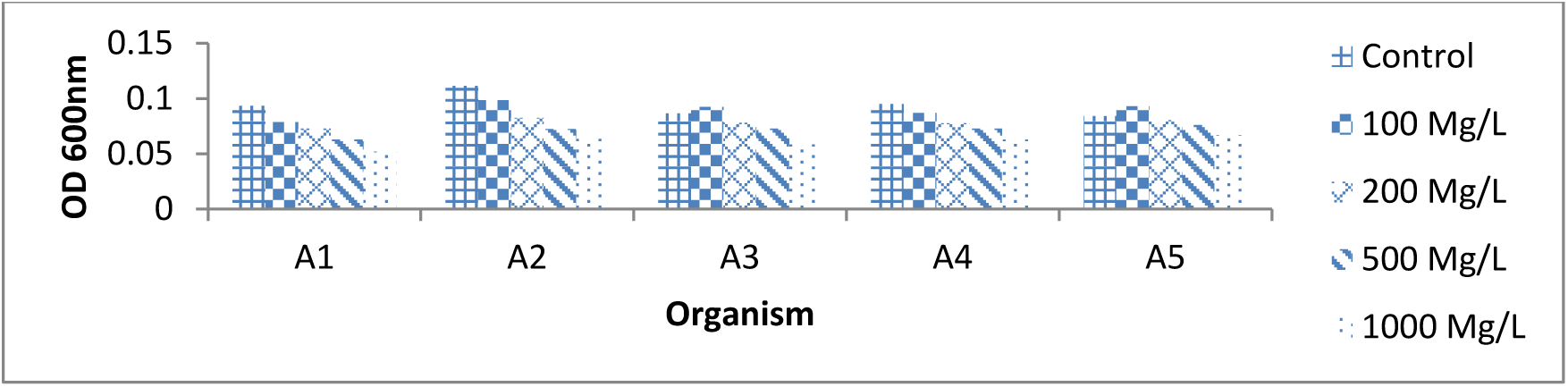
Response of different Ammonia Oxidizing Bacteria on different concentration of Lead within 120 hours.

**Figure 27:**
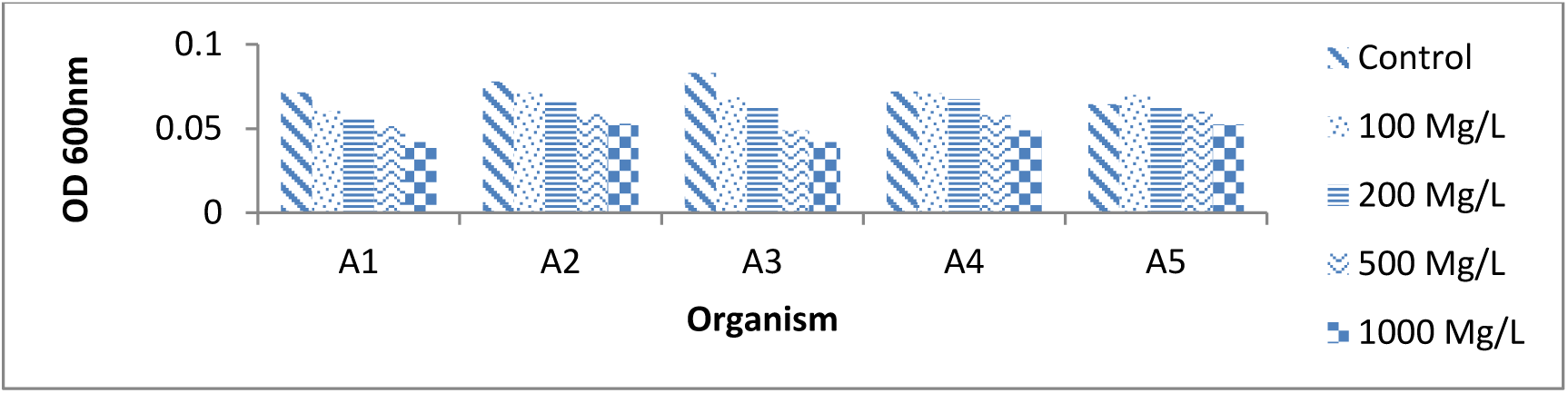
Response of different Ammonia Oxidizing Bacteria on different concentration of Lead within 168 hours.

### Tolerance of isolates to the cadmium

The results represented in Figures 28, 29. 30, 31 and 32 shows the level of tolerance for the different metal salt concentrations by the ammonia oxidizing bacteria isolates. Figure 28 shows the tolerance of the A1 isolates for different concentrations of Cd. The growth of A1 range from 0.0755 ± 0.006 to 0.0855± 0.001 within 72 hours, increase to 0.0885 ± 0.001 within 120hrs and decrease to 0.0805 ± 0.001 after 168 hours at 0 concentration. A1 growth range from 0.075 ± 0.0099 to 0.088 ± 0.008 within 72 hours, increase to 0.0849 ± 0.001 within 120hrs and decrease to 0.0635 ± 0.002 after 168 hours at 100ug/ml concentration. 0.0645 ± 0.006 to 0.0595 ± 0.006 within 72 hours; decrease to 0.052±0.014 within 120hrs and decrease to 0.025 ± 0.007 after 168 hours at 1000ug/ml concentration.

**Figure 28:**
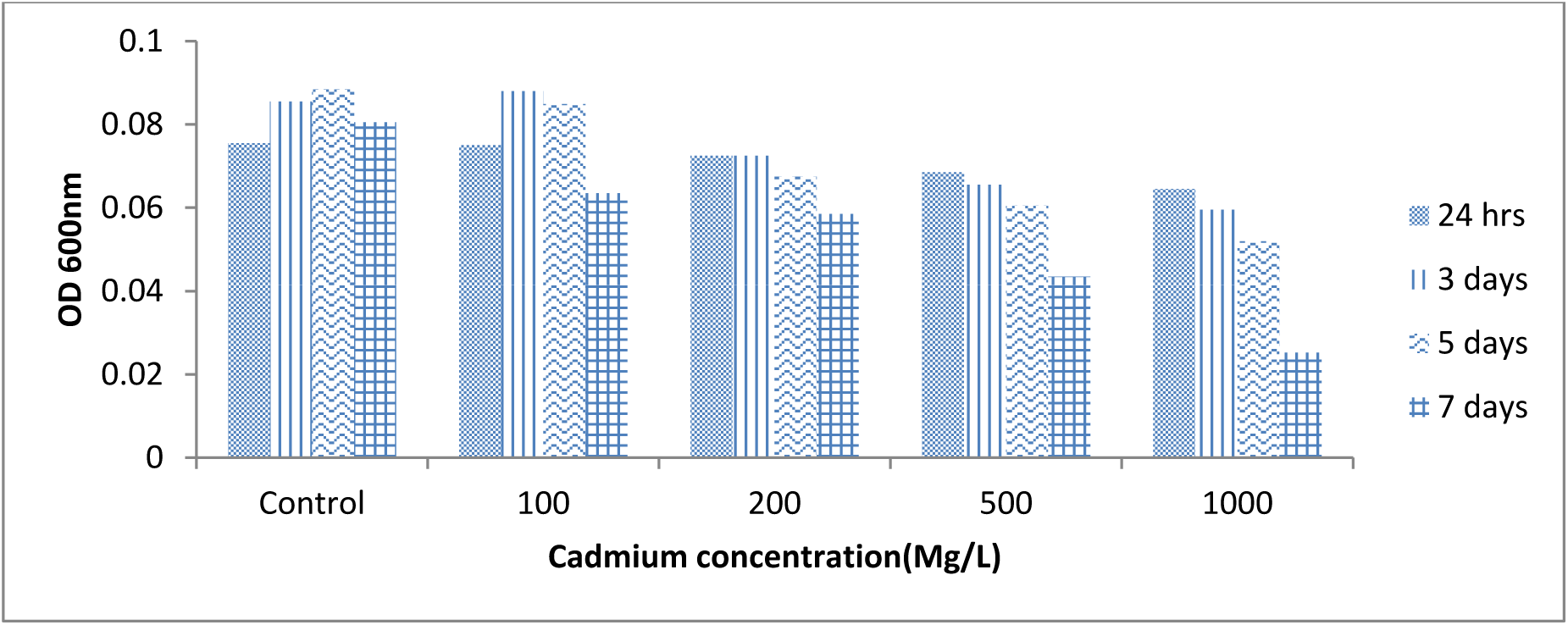
Tolerance of A1 to different cadmium concentration.

**Figure 29:**
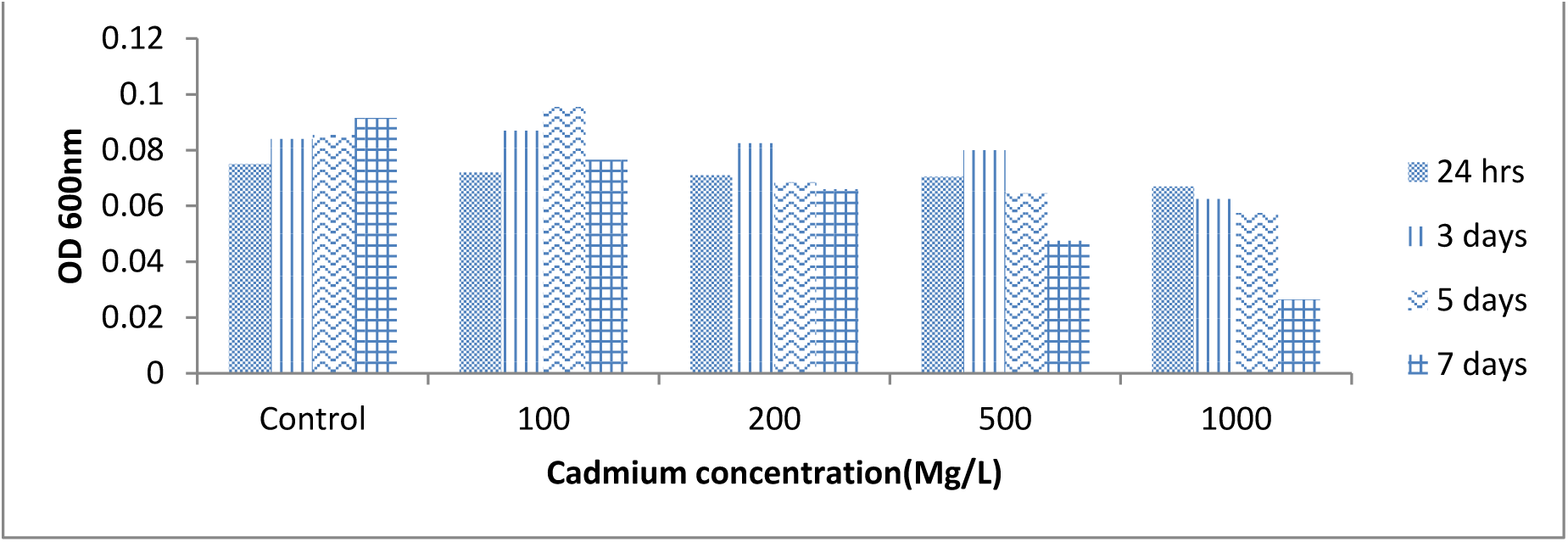
Tolerance of A2 to different cadmium concentration.

**Figure 30:**
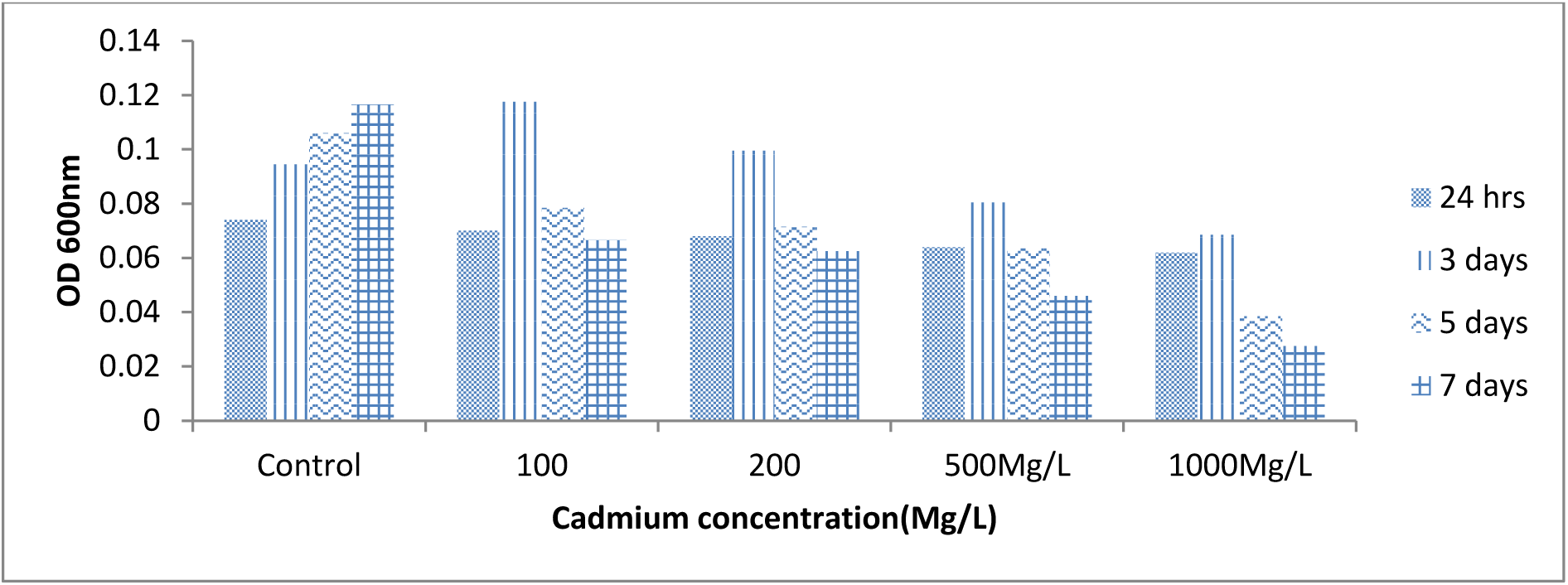
Tolerance of A3 to different cadmium concentration.

**Figure 31:**
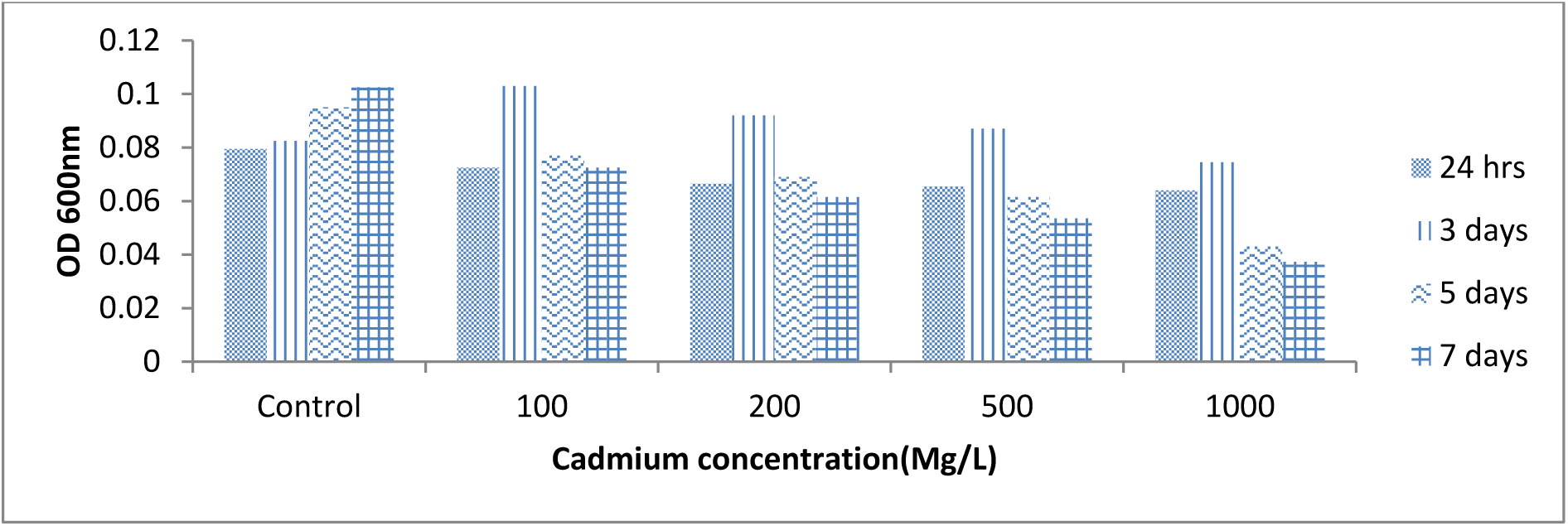
Tolerance of A4 to different cadmium concentration.

**Figure 32:**
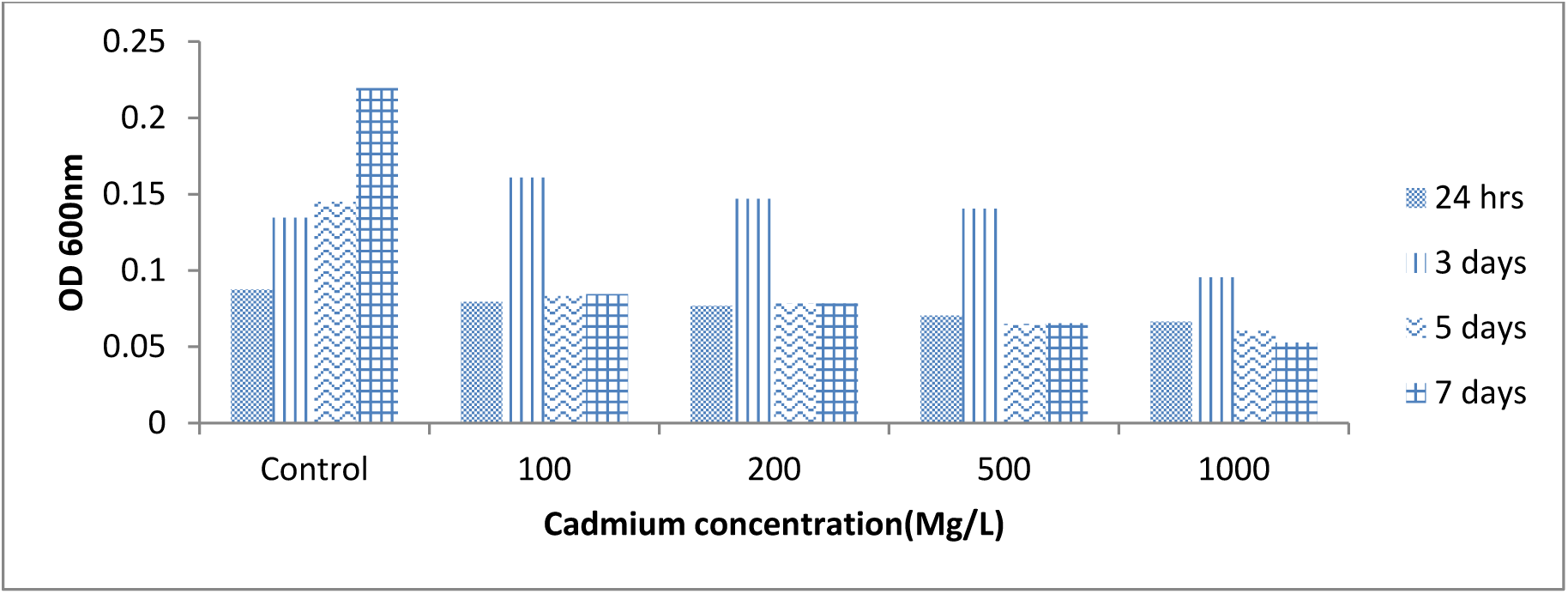
Tolerance of A5 to different cadmium concentration.

Figure 29 shows the tolerance of the A2 isolates for different concentrations of Cd. The growth of A2 range from 0.075 ± 0.008 to 0.084 ± 0.001 within 72 hours, increase to 0.0855 ± 0.001 within 120hrs and increase to 0.0915 ± 0.001 after 168 hours at 0 concentration. At 100ug/ml concentration A2 growth range from 0.072 ± .015 to 0.087 ± 0.008 within 72 hours, increase to 0.0955 ± 0.001 within 120hrs and decrease to 0.0765 ± 0.001 after 168 hours at 100ug/ml concentration. 0.067 ± .0006 to 0.0625 ± 0.002 within 72 hours; decrease to 0.0575 ± 0.001 within 120hrs and reduce to 0.0265 ± 0.002 after 168 hours

Figure 30 shows the tolerance of the A3 isolates for different concentrations of Cd. The growth of A3 range from 0.074 ± 0.014 to 0.0945 ±.0.002 within 72 hours, increase to 0.106 ± 0.003 within 120hrs and increase to 0.117 ± 0.015 after 168 hours at 0 concentration. A3 growth range from 0.07 ± 0.014 to 0.1175 ± 0.03 within 72 hours; decrease to 0.0785 ± 0.01 within 120hrs and decrease to 0.0655 ± 0.001 after 168 hours at 100ug/ml concentration. A3 growth range from 0.062 ± .0014 to 0.0685 ± 0.001 within 72 hours, decrease to 0.0385 ±0.001 within 120hrs and reduce to 0.0275 ± 0.001 after 168 hours at 1000ug/ml concentration.

Figure 31 shows the tolerance of the A4 isolates for different concentrations of Cd. The growth of A4 range from 0.0795 ± 0.005 to 0.0825 ± 0.002 within 72 hours, increase to 0.095 ±0.00 within 120hrs and increase to 0.103 ± 0.013 after 168 hours at 0 concentration. A4 growth range from 0.0725 ± 0.004 to 0.103 ± 0.011 within 72 hours; decrease to 0.077 ± 0.008 within 120hrs and decrease to 0.0725 ± 0.001 after 168 hours at 100ug/ml concentration. A4 growth range from 0.064 ± 0.008 to 0.0745 ± 0.001 within 72 hours, decrease to 0.043 ± 0.006 within 120hrs and reduce to 0.037 ± 0.002 after 168 hours at 1000ug/ml concentration.

Figure 32 shows the tolerance of the A5 isolates for different concentrations of Cd. The growth of A5 range from 0.0875 ± 0.002 to 0.1345 ± 0.002 within 72 hours, increase to 0.145 ± 0.001 within 120hrs and increase to 0.22 ± 0.02 after 168 hours at 0 concentration. A5 growth range from 0.0795 ± 0.001 to 0.161 ± 0.004 within 72 hours, decrease to 0.0833 ± 0.001 within 120hrs and increase to 0.0845 ± 0.006 after 168 hours at 100ug/ml concentration. A5 growth range from 0.067 ± 0.005 to 0.096 ± 0.02 within 72 hours; decrease to 0.061 ± 0.001 within 120hrs and decrease to 0.053 ± 0.00 after 168 hours at 1000ug/ml concentration.

The results represented in Figures 33, 34, 35 and 36 shows the level Response of different Ammonia Oxidizing Bacteria on different concentration of Cadmium within 24, 72, 120 and 168 hour respectively.

**Figure 33:**
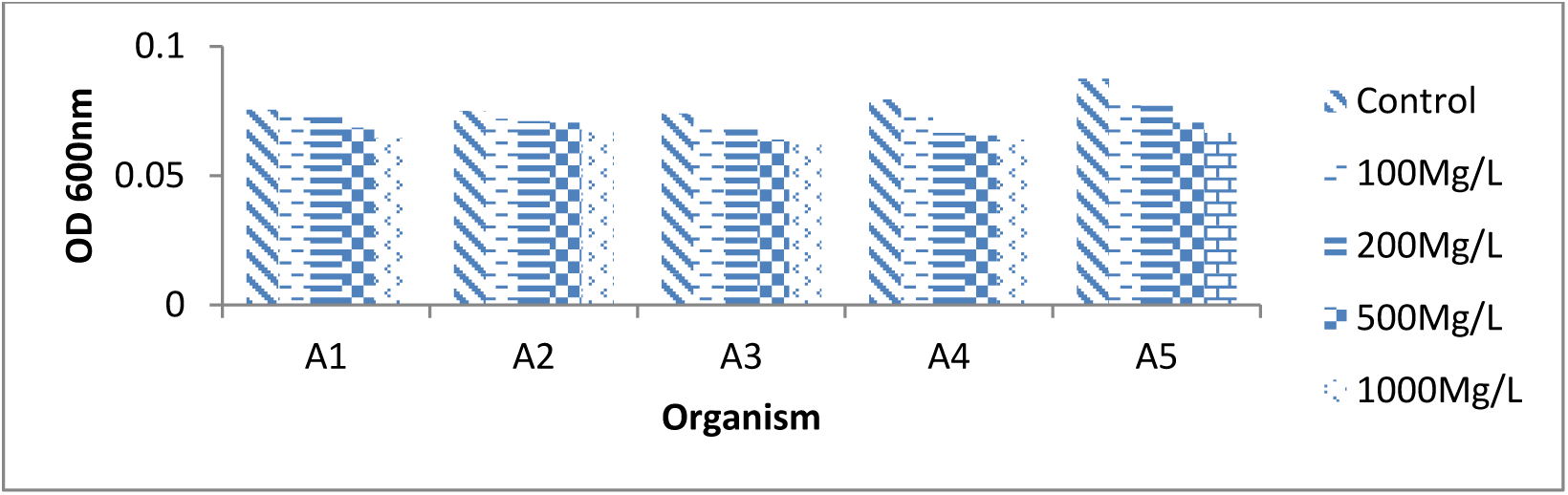
Response of different Ammonia Oxidizing Bacteria on different concentration of Cadmium within 24 hours.

**Figure 34:**
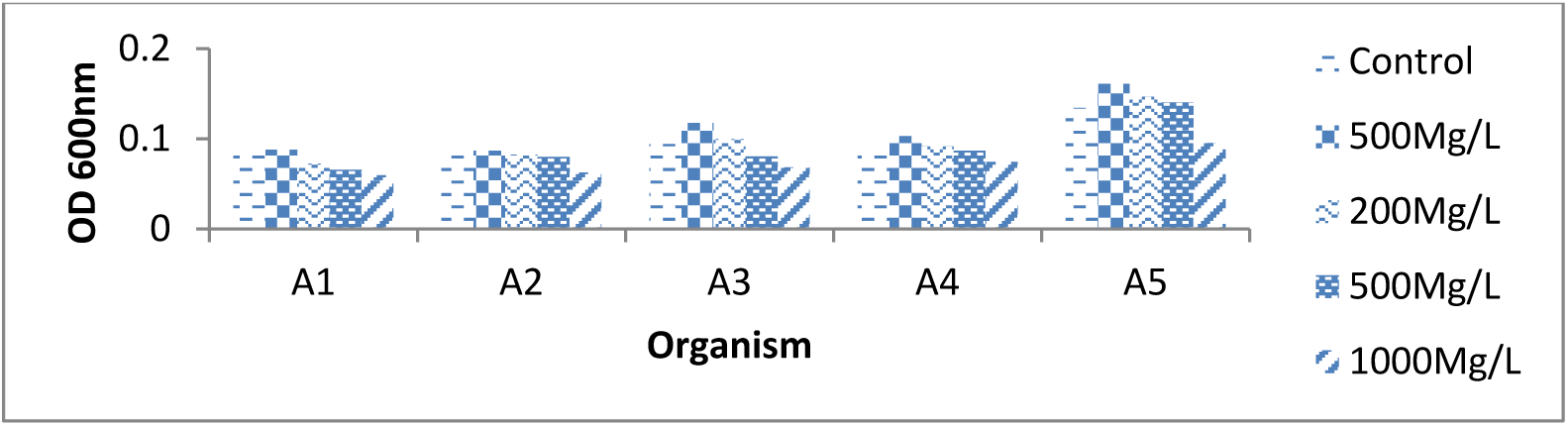
Response of different Ammonia Oxidizing Bacteria on different concentration of Cadmium within 72 hours.

**Figure 35:**
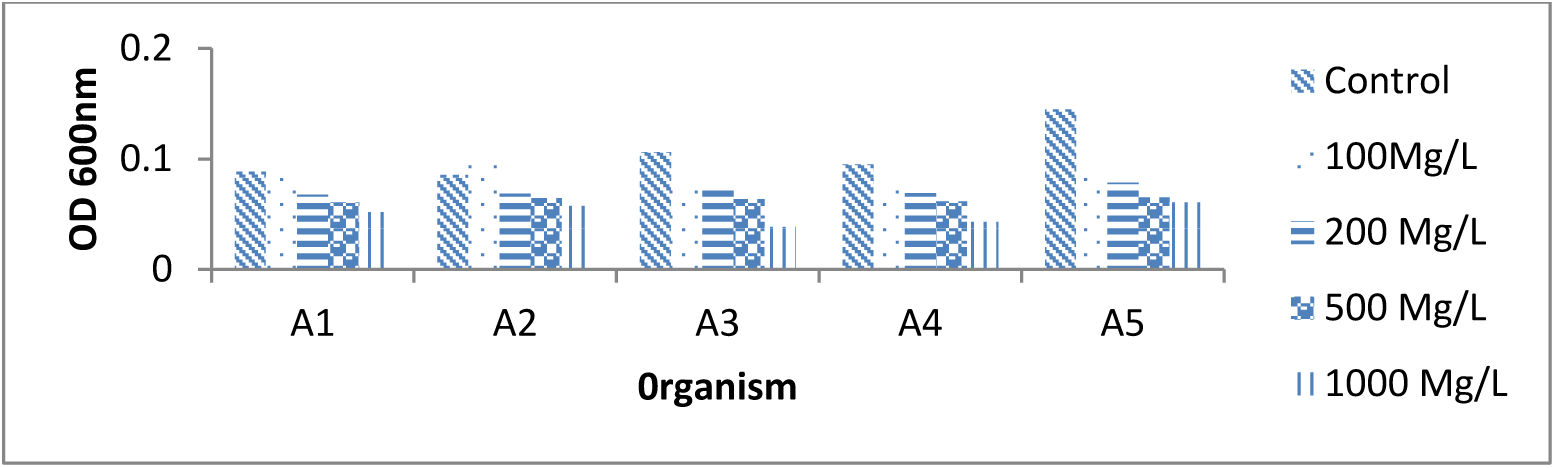
Response of different Ammonia Oxidizing Bacteria on different concentration of Cadmium within 120 hours.

**Figure 36:**
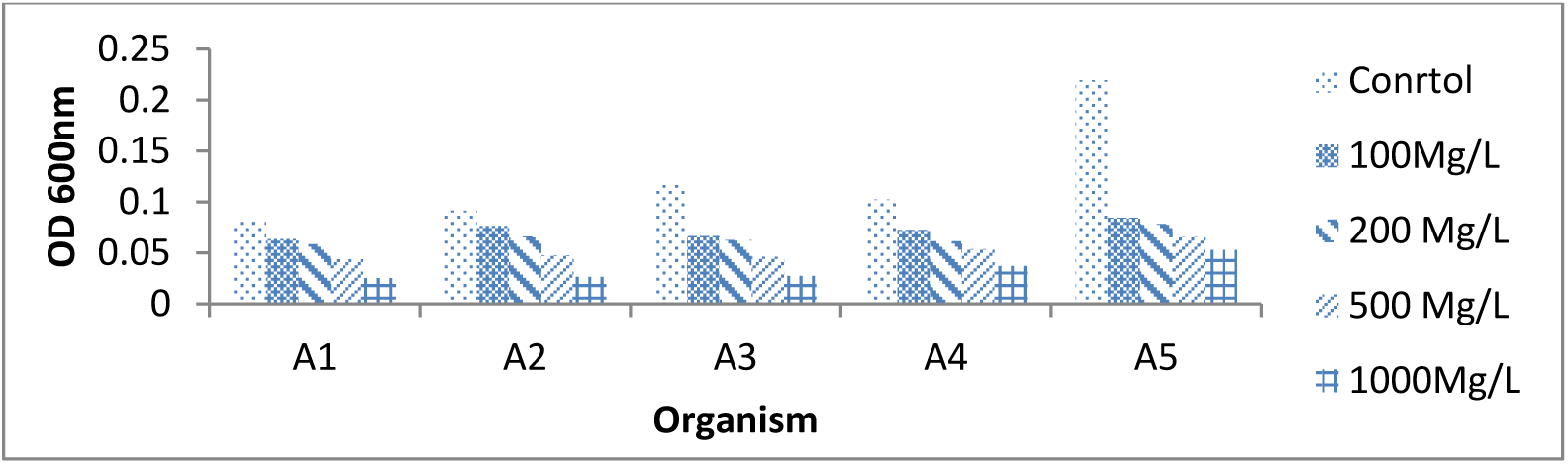
Response of different Ammonia Oxidizing Bacteria on different concentration of Cadmium within 168 hours.

## DISCUSSION

All the bacterial isolates show the ability to grow in Mineral salt agar medium incorporated with different heavy metal salt at different concentration (100ppm, 200ppm, 500ppm & 1000ppm). The cultures were incubated for 7 days and measured for optical density (at wavelength = 600 nm) in UV spectrophotometer. All bacteria showed high tendency to decrease optical density while increasing metal concentration in the medium when compare to control.

The growth of the Ammonia oxidizing organism increased successively throughout the period of five days and decrease at 7 days exposure time at 0(Control), 100 mg/L, 200 mg/L, 500 mg/L and 1000 mg/L concentration of metal salt. The highest growth was observed, for all the isolates, in the medium with no metal ion amendment, which served as the control, followed by 100 mg/L, 200 mg/L, 500 mg/L and 1000 mg/L respectively. Increased in the growth of ammonia oxidizing organism was observed throughout the period of five days and decrease mostly at 7 days exposure time at 0(Control), 100 mg/L, 200 mg/L. The sensitivity of the isolates to Salts of Cupper (Cu), Nickel (Ni), Cadmium (Cd), and Lead (Pb) where in the decreasing order A1> A3> A4 > A2> A5. All the isolates showed the high sensitivity to high concentration of 500 and 1000mg/L salts of Cupper (Cu), Nickel (Ni), Cadmium (Cd), and Lead (Pb). A2 and A5 exhibited the greatest ability to tolerate the metal salts than A1, A3 and A4.

Tolerance for the metal ions was dependent on concentration, time and the isolate tested (Siokwu and Anyanwu,2011). Khan *et al.* (2010); Subrahmanyam *et al.*,(2014); Wang *et al.*, (2017) reported that the toxicological effects of metals on soil microbial community structure and activities depend largely on the type and concentration of metal and incubation time. This finding revealed that when tolerance to the four metals was compared with respect to each bacteria isolate, it was evident that the bacteria were more sensitive to Cu^2+^ than the other three metals(Ni^2+^, Pb^2+^ and Cd^2+^), The sensitivity of the metal where in the order of Cu > Cd > Ni > Pb.

Microorganisms are very sensitive; they react quickly to any kind of changes (natural and anthropogenic) in the environment, and quickly adapt themselves to new conditions conditions including high metal concentrations. Microorganisms take heavy metals into the cell in significant amounts. This phenomenon leads to the intracellular accumulation of metal cations of the environment and is defined as bioaccumulation (Wolejko *et al.*, 2016). Some bacterial plasmids contain specific genes for resistance to toxic heavy metal ions (Liu *et al.*, 2018; Pacwa-Plociniczak *et al.*, 2018; Lukina *et al.*, 2016; Sharma, 2016), ability to produce siderophore, and ability to solubilize phosphate (biofertilizers) (Ibiene *et al.*, 2012; Gupta *et al.*, 2014). Some microorganisms can adjust their metabolic activity or community structure to adapt to the harmful shock loadings. Microorganisms play important role in stress environment and the derived ecosystem functions (Singh *et al.*, 2016a, b, c; Vimal *et al.*, 2017). Metals detoxification through resistance and tolerance, this resistance can be attributed to mechanisms of exclusion or tolerance (Klassen *et al.* 2000). These mechanisms stem from prior exposure of microorganisms to metals which enable them to develop the resistance and tolerance useful for biological treatment (Viti *et al.*, 2003; Sharma, 2016; Singh *et al.*, 2016a, b, c)

Nitrifying bacteria could be reduced by heavy metals and therefore considered as sensitive microbial process with regards to heavy metal stress (Smolders *et al*., 2001). Lee *et al.*, 2011; Liu *et al.*, 2010; Mertens *et al.*, 2009; Vasileiadis *et al.*, 2012 have particularly emphasized the response of ammonia oxidizing bacteria to heavy metals such as Zn, Cu and Hg. Frey *et al.*, 2008; Lee *et al.*, 2011; Vasileiadis *et al.*, 2012 have reported a sensitive response of the AOB community to heavy metals. Heavy metals can affect diversity of certain microbial communities and related soil processes (Haferburg and Kothe, 2007; Li *et al.* 2009a, b). Li *et al.* (2009a, b) reported that an increase in available Cu (0∼2400 mg kg^-1^) was paralleled by a concomitant decrease in AOB amoA copy numbers. Heavy metals used in greater amounts, result in metabolic disorders, suppress the growth of most plants and microorganisms (Ashraf and Ali, 2007, Zimmer *et al.*, 2016).

Markedly different responses of AOB communities to metal pollution stress have been observed Both metal-sensitive and metal-tolerant AOB populations have been observed in agricultural soils amended with metals (Stephen *et al.* 1999). Ferry reported that soil metal contamination did not decrease the abundance of AOB (Frey *et al.* 2008), while the research of Stefanowicz *et al.* (2008) and Qu *et al.* (2011) found that bacterial functional diversity was significantly decreased with increasing soil pollution. Bermudez *et al*. (2009) hypothesize that high organic matter contents of soil can bind metals and decrease their toxicity. Wu *et al.* 2011 investigated that there were no significant differences in the effect of metals on AOB abundance which suggested that metal concentration was not the main factor affecting the abundance of ammonia-oxidizing bacteria.

Moreover, Kris *et al.* (2005) had demonstrated that the nitrifying community displays a tolerance to long-term Zn stress (Rotthauwe *et al.* 1997). Liu *et al*. (2010) also found that there was no significant difference in the abundance of AOB among different concentrations of Hg.

## CONCLUSION

All the Ammonia Oxidizing bacteria were able to adapt and grow under various extreme conditions show a high level of tolerance for heavy metal tested. A2 and A5 exhibited the greatest ability to tolerate the metal salts than the others which makes the organism an attractive potential candidates for further investigations regarding their ability to remove heavy metal in bioremediation. It may be a good option for bioremediation of soil and waste since it is regarded as an eco-friendly and efficient. The understanding of microbial tolerance and adaptation to the presence of metal in the environment is critical in determining the management and potential long-term effect of that part of the environment receiving metal contamination.

## ACKNOWLEDGEMENTS

We thank the University of Nigeria Nsukka for giving us the opportunity to carry out the research and we also appreciate Rev. Fr. Prof. Vincent Nyoyoko, for funding the research.

